# A CRISPR homing screen finds a chloroquine resistance transporter-like protein of the *Plasmodium* oocyst essential for mosquito transmission of malaria

**DOI:** 10.1101/2024.06.02.597011

**Authors:** Arjun Balakrishnan, Mirjam Hunziker, Puja Tiwary, Vikash Pandey, David Drew, Oliver Billker

## Abstract

Genetic screens with barcoded *Plasmo*GEM vectors have identified thousands of *Plasmodium* gene functions in haploid blood stages, gametocytes and liver stages. However, the formation of diploid cells by fertilisation has hindered the use of genetic screens to investigate vector-parasite interactions during the mosquito stages of the parasite. In this study, we developed a scalable genetic system that uses barcoded gene targeting vectors equipped with a CRISPR-mediated homing mechanism to generate homozygous loss-of-function mutants to reveal gene functions in the functionally diploid life cycle stages. In this system, a knockout vector additionally expressing a gRNA for its target is integrated into one of the parental alleles and directs Cas9 to the intact allele after fertilisation, leading to its disruption. We find that this homing strategy is 90% effective in the oocyst, resulting in the generation of homozygous genotypes. A pilot screen reveals that PBANKA_0916000 encodes a chloroquine resistance transporter-like protein, CRTL, essential for oocyst growth and sporogony. The data point to an unexpected importance for the transmission of malaria of the poorly understood digestive vacuole of the oocyst that contains hemozoin crystals. The new screening strategy provides a method to discover systematically and at scale the essential malaria transmission genes whose first essential functions are after fertilisation in the bloodmeal, enabling their potential as targets for transmission-blocking interventions to be assessed.

## INTRODUCTION

Malaria is a vector-borne disease caused by *Plasmodium* parasites and transmitted by female *Anopheles* mosquitoes, primarily resulting in fatalities among children under the age of 5. Vector control strategies, such as the use of long-lasting insecticide-treated nets and indoor residual spraying have averted 68% of malaria cases between 2000 to 2015^1^ but the emergence of insecticide-resistant mosquitoes and drug-resistant parasites have contributed to a reversal of this positive trend, resulting in a death toll of 608,000 as of 2022 (WHO report 2022). These data underscore the necessity for novel strategies to combat malaria transmission.

In an infected host, haploid malaria parasites replicate asexually within erythrocytes. While haploid forms cause the disease, transmission is initiated by sexual precursor stages, the gametocytes when a female *Anopheles* mosquito ingests an infectious blood meal. In response to environmental clues gametocytes in the blood bolus mature rapidly into haploid gametes, which fertilize to generate a diploid zygote. The zygote replicates its genome and undergoes meiosis to produce a tetraploid ookinete in which all four genomic products of meiosis persist within the same nuclear envelope. Motile ookinetes traverse the mosquito midgut epithelium and differentiate into oocysts. Over a period of 2 weeks in *P. berghei*, these cysts grow under the basal lamina and undergo endomitotic replication to generate and release infectious sporozoites, which invade the salivary glands of the mosquito. The significant population bottleneck posed by the transition from ookinetes to oocyst makes these stages vulnerable to transmission-blocking interventions, such as vaccines and anti-malarial drugs^2–4^. Nonetheless, the oocyst remains one of the least understood life-cycle stages of *Plasmodium*^5^.

Genetic screens, using barcoded knockout vectors of the *Plasmo*GEM (*Plasmodium* Genetic Modification) resource have revealed gene essentiality at different haploid life-cycle stages of the rodent malaria parasite *P. berghei*^6–10^. *Plasmo*GEM mutants can be screened simultaneously in pools containing dozens to many hundreds of barcoded mutants, each disrupted in a different gene. Although barcoded pools have been transmitted through mosquitoes to identify developmental blocks at the subsequent liver stage, such experiments have not been effective at revealing oocyst gene functions^7^ because all four products of meiosis persist within a single nuclear envelope and contribute to the proteome of the growing cyst. Once both parental genomes are active after fertilization, loss-of-function mutations that these tetraploid cells inherit from only one of the parents are often functionally complemented by the intact allele inherited from the other parent^7^. Genomes only segregate days or weeks later, when individual haploid sporozoites form within the oocyst. Most loss-of-function alleles will only reveal a phenotype days after that, depending on when any mRNA or protein that the sporozoite has inherited from the cyst will have turned over.

Here, we show that in the functionally diploid stages of *P. berghei*, gene functions can nevertheless be identified simultaneously and at scale by equipping barcoded knockout vectors with a guide RNA (*gRNA*)-based homing system. When integrated into one parental genome, a Cas9 endonuclease in the zygote can be targeted to the intact target locus of the other parental genome. In a pilot screen of 21 vectors, we demonstrate that gRNA homing is efficient enough to reveal new gene functions after fertilisation. We discover a structural homolog of the chloroquine resistance transporter (CRT), describe its importance for oocyst maturation and its localization to a putative digestive vacuole compartment in the oocyst.This study unveils a new homing-*gRNA*-based genetic screening approach in *Plasmodium* that can shed light on post-zygotic gene functions and biology of the oocyst, potentially identifying new targets for blocking parasite transmission.

## RESULTS

### Single sex *P. berghei* lines expressing Cas9-BFP are fertile

Sex in haploid malaria parasites is not chromosomally determined, and parasite clones produce both male and female gametocytes. We reasoned that if we could genetically modify lines to produce gametes of only one or the other sex, these could be engineered further, such that in a genetic cross each sex would separately deliver either a Cas9 endonuclease or a guide RNA to the zygote and cause a double strand break in a target gene. Since *Plasmodium* parasites lack canonical non-homologous end joining^11^, repair of double strand breaks is always homology driven. We therefore further reasoned that a disrupted target allele not recognised by the guide and carried by one of the parental lines would serve as repair template, leading to the disruption being copied from one parental genome to the other, generating a homozygous KO (Fig. 1).

**Figure 1.**
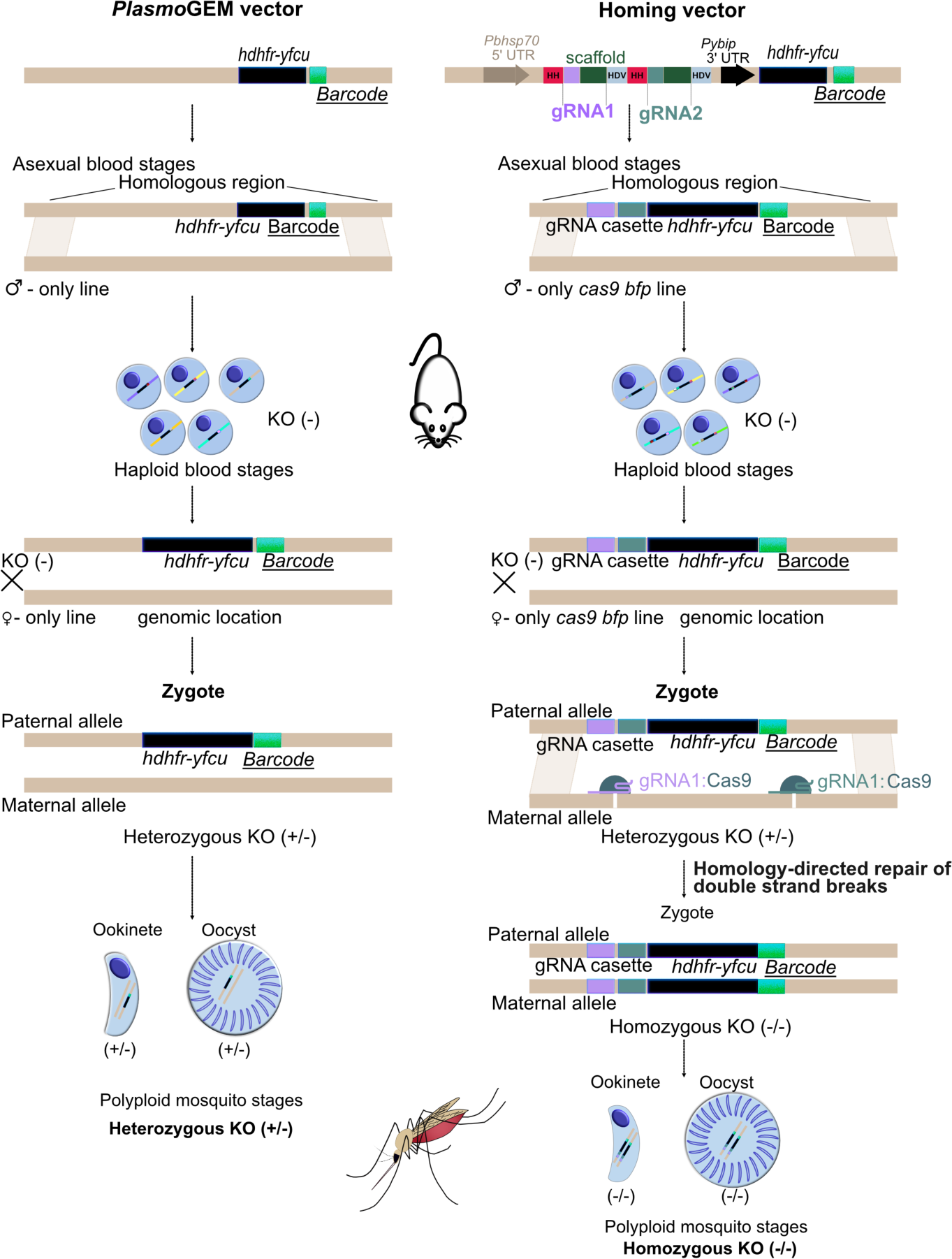
Schematic representation of the homing strategy. To generate homozygous knock-out ookinetes and oocysts (KO) after pooled transfection of modified *Plasmo*GEM vectors a gRNA and Cas9 are brought together in the zygote at the point of fertilisation. The disrupted allele in one parental genome is presumed to serve as repair template for a Cas9-mediated double strand break in the other parental genome.

To test this concept, we first created lines making only male or only female gametocytes by disrupting *female development 1* (*fd1*) or *male development 4* (*md4*), respectively (Fig. S1A), genes known to have sex-specific developmental functions early in gametocytogenesis^6^. These single-sex lines differed from those we recently used to screen for sex-specific fertility genes^10^ in that they did not express GFP, and in that after negative selection against the resistance marker, an expression cassette for Cas9 fused to a blue fluorescent protein (BFP) was left behind in the disrupted *fd1* or *md4* locus under the control of an *hsp70* “constitutive” promoter (Fig. S1A-C). Wild type parasites were eliminated by fluorescence-activated cell (FACS) on BFP followed by limiting dilution. The resulting clones expressed a flag-epitope-tagged Cas9-BFP fusion protein of the expected size (Fig. S1D), which was present in the nucleus of schizonts and, when lines were crossed, in zygotes and ookinetes (Fig. S1E, F).

Clonal *md4^-^cas9-bfp* (female-only) parasites produced female gametes expressing the P28 surface marker (Fig. S1F), did not produce male gametes (Fig. S1G) and gave rise to ookinetes only when co-cultured with the *fd1^-^cas9-bfp* (male-only), which did release male gamete, as expected (Fig. S1G). Also as expected, these single-sex lines produced no oocysts when transmitted individually, but oocyst formation was restored when mosquitoes were fed on mice co-infected with both lines (Fig. S1H), although not to the same level as wild type (Fig. S1H). This was not due to crosses *per se* being less efficient at producing oocysts, because two similar single sex lines not expressing Cas9^10^ were fully fertile when crossed. This led us to ask if Cas9 expression is toxic, but a line expressing Cas9 from the same control regions but inserted into the *p230p* locus made large numbers of oocysts (Fig. S1H). Further crosses between the different lines showed that a moderate reduction in oocyst numbers was associated specifically with the female-only *md4*^-^*cas9* clone, most likely due to a heritable variation in that clone that is unrelated to the expression of Cas9.

### Cas9-mediated genome editing after fertilisation is efficient

To quantitate the efficiency of Cas9-mediated genome editing after fertilisation at the level of individual oocysts, we designed a colour swapping experiment. This involved constructing complementary single-sex lines in which the endogenous MyoA protein was either fused to mCherry or to GFP (Fig. 2A and Fig. S2). MyoA is abundantly expressed during sporogony and tolerates c-terminal tagging^12^. We hypothesised that crossing single-sex lines expressing MyoA-mCherry and MyoA-GFP, respectively, should make yellow oocysts. However, if the latter locus additionally expressed one or more gRNAs targeting Cas9 specifically to the *mCherry* locus (Fig. 2A), homology-mediated repair of a double-strand break would use the complementary GFP-containing allele as repair template, turning oocysts green.

**Figure 2.**
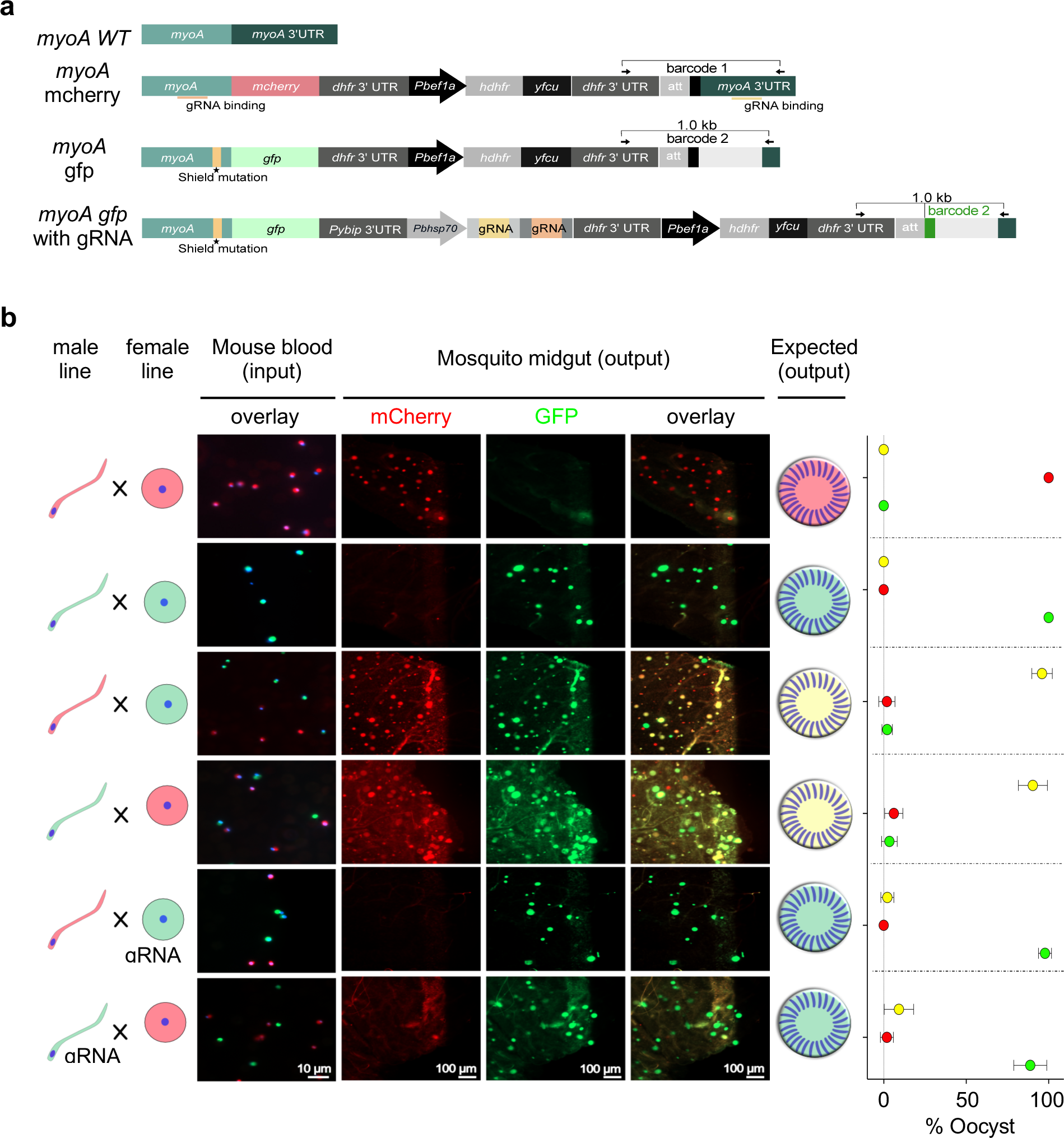
Colour-swapping of tagged MyoA variants as proof-of-principle for homing post fertilization. (a) Schematics of *myoA* tagging vectors. (b) Representative microscopic images of mouse blood (input) and oocysts on dissected mosquito midguts 12 days post infection (output). Mouse blood was stained with Hoechst (blue) before imaging. Graphs on the right show relative abundance of differently coloured oocysts from 25-50 mosquitoes analysed from two replicate transmission experiments per condition, each carrying at least 10 oocysts. Error bars show standard deviations of the means of all mosquitoes.

Consistent with this concept, directional crosses between red and green parents produced yellow oocysts, unless either the female or the male gamete provided gRNAs targeting the mCherry locus (Fig. 2B; all crosses also delivered Cas9 to the zygote). Since *md4* and *fd1* mutants do not undergo selfing, we were surprised to see 1-3% of oocysts in these crosses express only one parental allele, indicating that some zygotes may obtain an incomplete genome from a parent or that occasionally not all four products of meiosis survive or replicate in the oocyst. Our data suggest homing happens with 97% efficiency when the gRNAs are expressed from the female gamete, but this is reduced to 89% if the male carries the guides (Fig 2C, Table S1). This colour swapping experiment demonstrates that we have developed a CRISPR method where the phenotype of the oocyst is dominated by only one parental allele, in this case *myoa-gfp*, irrespective of the route of inheritance.

### Homing is scalable to screen for new post-fertilisation phenotypes

To ask if Cas9-mediated homing could reveal new loss-of-function phenotypes when pools of mutants were transmitted simultaneously, we inserted gRNA expression cassettes into each of 21 gene knockout vectors from the *PlasmoGEM* resource, using recombinase mediated engineering^13,14^ (Fig. S3). The resulting vectors used the CRISPR ribozyme-guide-ribozyme (RGR) strategy to express two *sgRNAs* from an RNA polymerase II promoter^15^ and were additionally equipped with the gene-specific DNA barcodes used routinely in *Plasmo*GEM screens^16^. Target genes were selected to be redundant in the asexual blood stages^9^, and to produce a range of known phenotypes to benchmark the system. We also included seven genes with known or likely functions in the functionally diploid mosquito stages, as suggested by their transcription patterns^17^, and three liver stage essential genes known not to have a role in oocysts to serve as reference points (Table S2).

We transfected *fd1^-^cas9* parasites either with a pool of the original *Plasmo*GEM vectors or with a pool of the corresponding homing vectors (Fig. 3A) and counted barcodes from the transfected vector pool (barcode input 1). Mutants selected with pyrimethamine (input 2) were then used to co-infect mice with another line that served as donor for fertile female gametocytes. Female *Anopheles stephensi* mosquitoes were then allowed to feed on six infected mice per pool. For each of two biological replicates, infected midguts of 80-100 mosquitoes per pool were dissected 12 days later, and barcodes counted in gDNA from the oocysts (barcode output) were compared to input 2, i.e. the barcode counts from the mice used to infect them (Fig. 3A).

**Figure 3.**
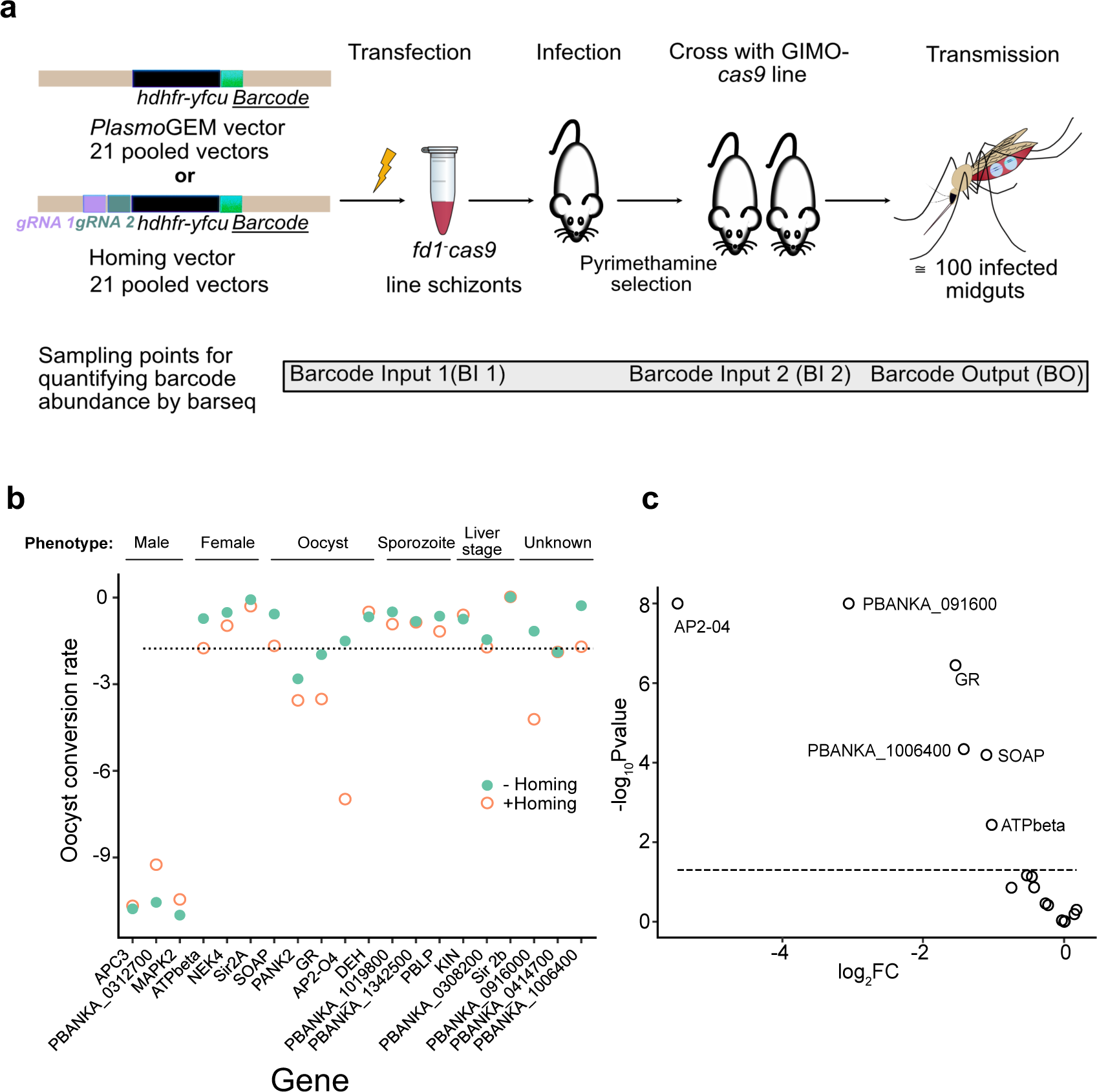
Pilot screen using CRISPR homing. **(a)** Schematic illustration of screen design. The male-only Cas9-expressing line was transfected with either homing or standard *Plasmo*GEM vectors. Transfected parasites were selected using pyrimethamine and then used to infect a new mouse together with a line providing female gametocytes. Sampling points for barcode counting are indicated. **(b)** Oocyst conversion rates derived from the relative abundances of barcodes in BO over BI 2 samples. Known phenotypes shown above. (**c)** Volcano plot showing the impact of homing as log2fold changes in relative conversion rates to oocysts. Each circle represents a gene. Effect sizes are plotted against the -log_10_ p-values calculated using a Z-test. The dotted line shows a p-value of 0.05. Male-specific genes were omitted from this analysis since their barcodes were lost from the pools irrespective of homing.

First, we asked if any *Plasmo*GEM homing vectors were underrepresented in the blood stages when compared to their corresponding non-homing version (Fig. S3B, C; Table S3). While most homing and non-homing vectors were equally abundant at the point of transfection (Fig. S3B), one homing vector targeting glycerol-3-phosphate dehydrogenase (G3PDH) was underrepresented after drug selection of the transfected parasites (Fig. S3C). Since G3PDH is dispensable in the blood stages, we concluded that the homing vector targeting G3PDH has either lost its functionality during recombineering or that one of the gRNAs has an off-target effect. This mutant was therefore removed from further analysis.

To find new gene functions in the oocyst, we looked for cases where the presence of gRNAs in the KO vector affected the conversion rate of asexual blood stages into oocysts (Fig. 3B, Table S3). The oocyst conversion rate was first normalised to genes with liver-specific functions known to be dispensable at both blood and oocyst stage. Since a male-only line had been mutagenised, control mutants with known defects in male fertility, such as mitogen activated protein kinase 2 (ref^18^), were unsurprisingly depleted among the oocysts, irrespective of the homing capability of the vector (Fig. 3B). In contrast, for other genes homing vectors produced marked reductions in oocyst barcodes when compared with their non-homing equivalents (Fig. 3C). For the transcription factor *ap2-o4,* for instance, gRNAs caused a 64-fold reduction, consistent with its known essential role between ookinete formation and the oocyst^19^. Screen data for glutathione reductase (*gr*) and the secreted ookinete adhesive protein (*soap*) also reproduced known phenotypes of cloned mutants^20,21^ which were not previously visible in *Plasmo*GEM screens. On the other hand, the screen did not reveal a phenotype for 3-hydroxy acyl-CoA dehydratase (*deh)*, possibly because the late oocyst degeneration observed in a cloned mutant in this gene^22^ is not accompanied by a sufficient reduction in genome copies. Two of three oocyst-expressed genes of unknown function, PBANKA_0916000 and PBANKA_1006400, showed a marked homing effect warranting further investigation.

### PBANKA_0916000 encodes an essential digestive vacuole protein related to the chloroquine resistance transporter

How mutants behave in the homing screen may be confounded by factors other than gene function, such as the efficiency of gRNAs, how much transcript or protein a zygote inherits from the female parent and how fast these turn over. To validate the result of the pilot screen we studied PBANKA_0916000 a poorly understood gene with a new phenotype, which we find encodes a chloroquine resistance transporter like (CRTL) protein.

To understand the role of CRTL in mosquito stages, we first fused a c-terminal triple HA epitope tag to the endogenous gene (Fig. S4A, B) to monitor expression and localisation of the protein. Western blot analysis suggests absence of CRTL from asexual blood stages, but a full-length (132 kDa) fusion protein was detected in ookinetes (Fig. S4D), which is consistent with mRNA abundance peaking in ookinetes according to the Malaria Cell Atlas^17^. Fluorescence microscopy confirmed this observation and showed a small number of weakly labelled foci first appearing in the ookinete cytosol (Fig. S4E). A time course of oocyst development detected CRTL-HA from day 2, where the label colocalised with highly refractile hemozoin crystals (Fig. 4). Hemozoin in the oocyst is thought to originate from the macrogametocyte which, like all erythrocytic stages, detoxifies heme moieties from haemoglobin digestion by aggregating them inside the digestive vacuole in crystalline form. Hemozoin is carried over into the oocyst where it apparently continues to reside inside the remnants of a DV-like compartment labelled by CRTL-HA. HA-positive granules are spread throughout the cytoplasm on days 2 and 4 of oocyst development. From day 6 they form aggregates, which do not increase in size as the oocyst grows. Around days 8-10 these labelled structures become forced into the narrowing spaces between growing sporogonic islands, where they continue to colocalise with the refractile pigment crystals (Fig. 4) whose arrangement into lines or crosses is a hallmark of the oocyst at this stage. When oocysts sporulate, neither HA-tagged protein nor hemozoin crystals get incorporated into sporozoites (not shown). These results are consistent with the tagged protein residing in the DV remnants surrounding hemozoin crystals.

**Figure 4.**
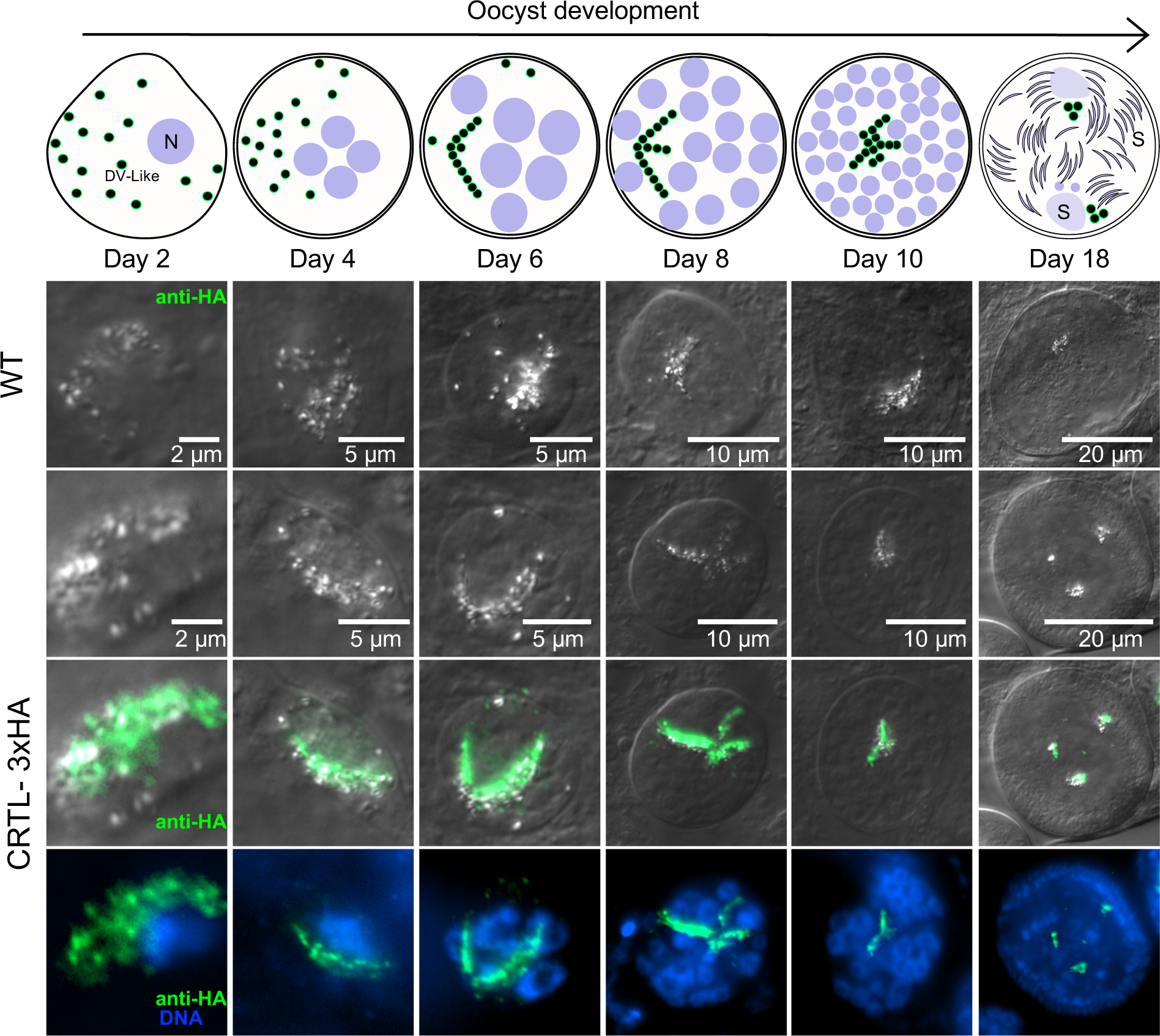
Immunofluorescence micrographs localizing CRTL-3xHA. Midguts infected with wild type parasites and parasites carrying a 3xHA tag on CRTL were dissected and fixed on different days post-infection. Immunostaining against HA shown in green, Hoechst-stained DNA shown in blue. The schematic above illustrates the digestive vacuole like structures (DV-like), nuclei (N), sporoblast areas (SB) and sporozoites (SP) at different stages of oocyst development.

We created a clonal *crtl* KO line in an mCherry expressing background (Fig. S4A, C) and compared its growth to a GFP expressing WT (Pb Bergreen) clone. Although CRTL was first detected in ookinetes, mutant gametocytes converted to ookinetes in culture at the same rate as wild type (Fig. S5B). Consistent with this, there was no difference in the number of oocysts between WT and mutant on day 10 or 14 (Fig. 5A; Table S1), but mutant oocysts were smaller on both days (Fig. S5A). In a time-course experiment, mutant oocysts stopped growing from day 8 (Fig. 5B, C). By day 18, WT oocysts had grown to more than twice the diameter of mutant oocysts (Fig. 5B; Table S1).

**Figure 5.**
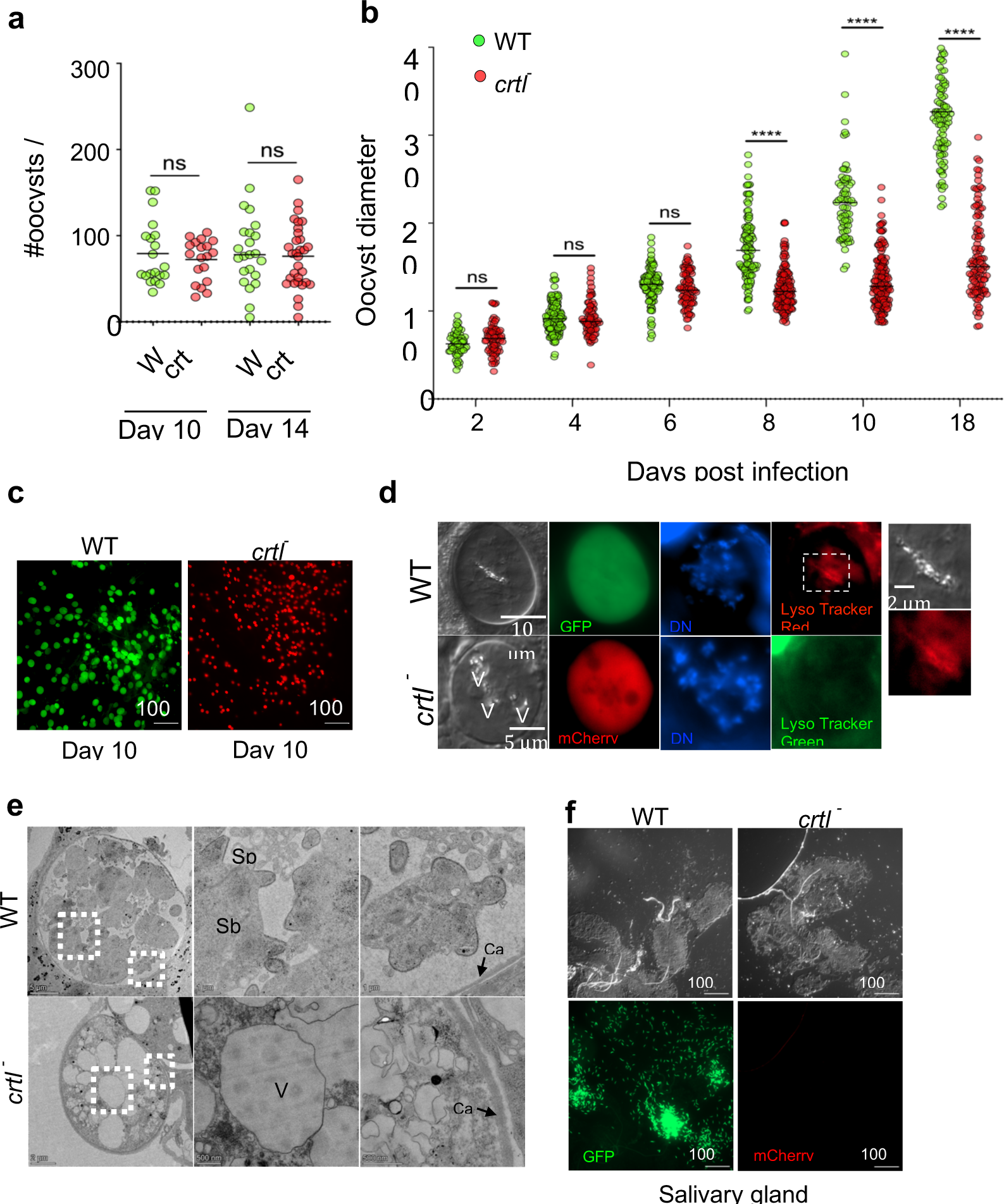
Characterisation of a cloned CRTL mutant. **(a)** Oocyst numbers on 25 dissected guts from infections with Pb Bergreen (WT) or *Pb mcherry crtl^-^* parasites. The data are representative of two infections. (ns, not signifnicant in unpaired T-test). **(b)** Growth kinetics of oocysts from 30 infected mosquitoes per time points (ns, not significant; ****P < 0.0001 in unpaired T-test). **(c)** Representative images illustrating Lysotracker staining oocyst size differences. **(d)** Representative images of day 8 oocysts highlighting acidic compartments with LysoTracker staining. The boxed area is enlarged on the right to show DV-like acidic compartment in a WT oocyst. V, vacuolar structures that appear in *crtl^-^* oocysts. **(e)** Representative transmission electron (TEM) micrographs of day 10 oocysts. Boxed areas are enlarged to show sporoblast (Sb), capsule (Cap) and sporozoite (Sp) in WT oocyst. crtl^-^ oocysts are smaller, lack sporoblast and have enlarged vacuolar structures (V) despite having an intact capsule (cap). **(f)** Representative images of a set of salivary glands of WT or *crtl^-^*infected mosquitoes 20 days post-infection.

On the day when growth was first reduced, mutant oocysts had commenced DNA replication like wild type, but were characterised by round vacuoles which excluded cytosolic mCherry protein (Fig. 5D). By day 10 virtually all mutant oocysts appeared vacuolated (Fig. S5C). Time lapse imaging showed Brownian movement of pigment crystals inside the vacuoles, suggesting vacuolation results from swelling of DVs (Fig. S6, Supplementary Video 1). In wild type oocysts pigment crystals colocalized with lysotracker, consistent with them residing inside an acidic compartment (Fig. S6). In contrast vacuoles of mutant oocysts did not stain with lysotracker, suggesting a swelling of the DV is accompanied by a loss of acidification. In contrast, ookinetes, which we showed not to require CRTL, possess lysotracker-positive acidic compartments also in the mutant (Fig. S5D).

Transmission electron microscopy performed on 10-day-old oocysts confirmed that the smaller mutant parasites contained vacuoles of around 1 µm diameter (Fig. 5E). Unlike wild type oocysts, where sporoblasts (Sb) and forming sporozoites (Sp) were visible on day 10, knock-out oocysts lacked signs of sporulation. Mutant parasites also failed to produce salivary gland sporozoites compared to wild type on day 20 (Fig. 5F). Mosquitoes infected with mutant were unable to infect mice by bite, showing the gene is essential for transmission (Fig. S5E).

An AlphaFold 2 model^23^ predicts that PbCRTL has the drug-metabolite-transporter (DMT) fold^24^, which in vertebrates is principally utilized by transporters for nucleotide-sugars and nucleotides^25^ (Fig. 6A). In lower organisms, however, bacterial proteins (e.g., YddG, YdeD, and RhtA) have been shown to serve as exporters of amino acids and their derivatives^26–28^.

**Figure 6.**
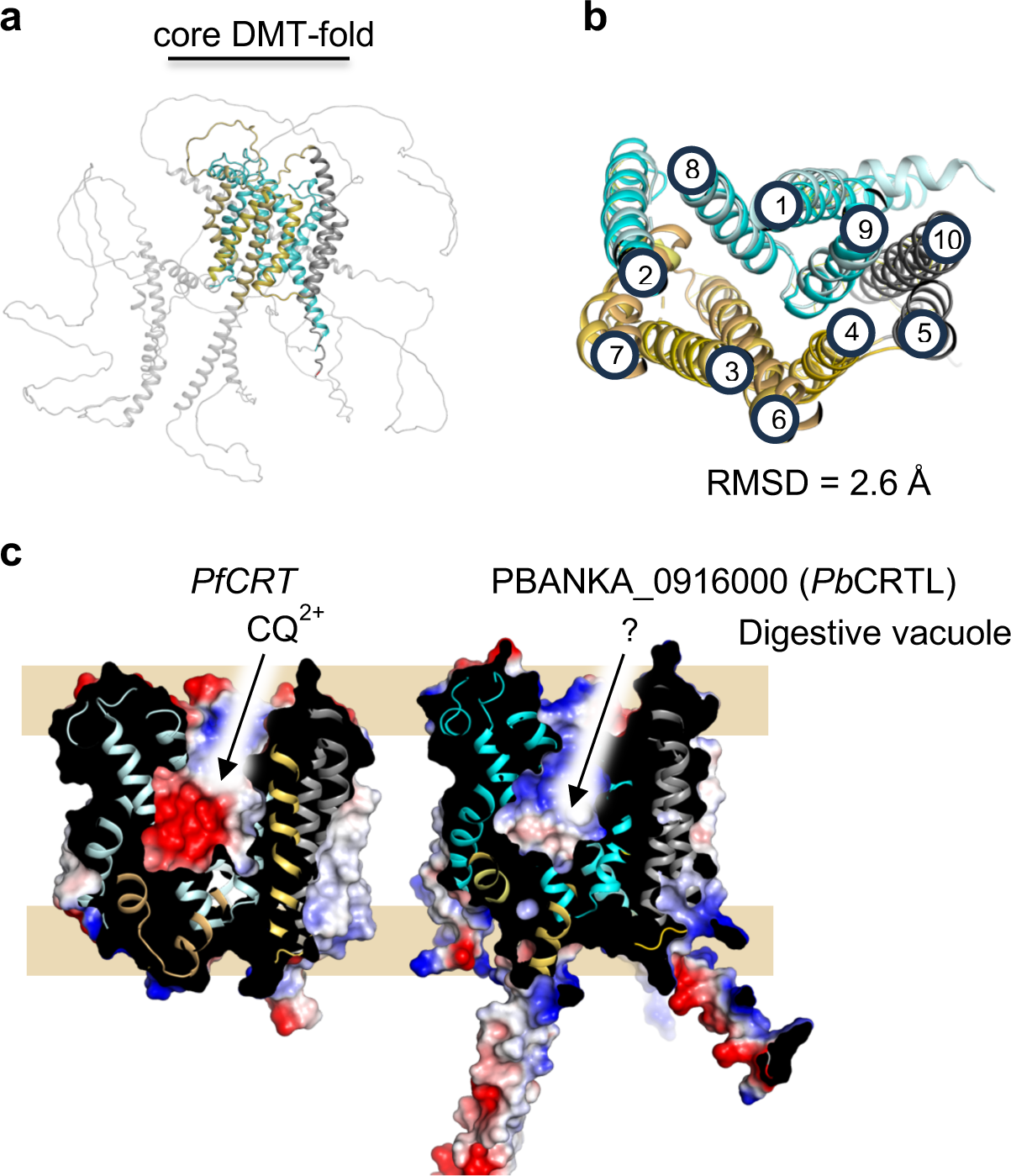
CRTL structural modelling. **(a)** AlphaFold2 (AF2) predicted structure of *Pb*CRTL. The core of the Drug Metabolite Transporter (DMT) fold is colored based on the bundle domains (orange and cyan) that alternate access using a rocker-switch mechanism. Regions of predicted disorder and extramembrane helices are shown in light-grey. **(b)** Cartoon representation of a structural superimposition of *Pb*CRTL and *Pf*CRT (RMSD of 2.6 Å) viewed from the digestive vacuole lumen. The two bundles of the DMT-fold that alternate access are colored in cyan and yellow for *Pb*CRTL and light-blue and sand for *Pf*CRT, respectively. TM5 and TM10 in grey can form an oligomeric interfaces in some members and show larger variation. (c) An electrostatic potential surface representation of PfCRT and *Pb*CRTL protein highlighting the charge differences in the substrate binding cavity; colored blue to red, for positive to negative charge.

Using FoldSeek^29^, the closest experimental structure to PBANKA_0916000 is the Chloroquine Resistance Transporter from *Plasmodium falciparum* (*Pf*CRT)^30^. Indeed, a superimposition of the AF2 prediction and *Pf*CRT shows a high degree of structural similarity with root mean square deviation of 2.9 Å^24^ (Fig. 6B). In addition to the core DMT-fold the *Pb*CRTL protein has extensive loops and tails with intrinsically disordered domains (IDRs). Such extra-membranous regions are not required for transport, and it is most probable that CRTL forms a complex with other proteins that regulate its activity. OrthoDB identifies CRTL homologs in all *Plasmodium* species, but not in other Apicomplexa. An alignment of these homologs using ClustalW revealed high sequence similarity within the conserved core of the putative transporter (Fig. S7, S8).

## DISCUSSION

The biology of *Plasmodium* ookinetes and oocysts, and their interactions with the vector remain poorly understood, although these stages determine vectorial capacity and the reproductive rate of malaria. In this study we demonstrate that a *gRNA*-based homing system makes functionally diploid mosquito stages tractable for the type of genetic screening approaches that have revealed thousands of gene functions in haploid stages ^6–10,31^. In a proof-of-concept colour swapping experiment we found homing was 89% effective when the male gamete provided the gRNA and reached 97% when the gRNA entered the zygote with the female gamete. This difference may be due to the *hsp70* promoter being less active in males, or to the compacted male nucleus delivering less Cas9-gRNA riboprotein complex to the zygote. We cannot say precisely when after fertilisation Cas9 targets the *mCherry* gene, but if transcriptional activation of the male genome is required, this may also reduce efficiency because some paternally inherited genes remain transcriptionally silent for varying periods after fertilisation^32^. Whatever the precise timing and mechanism, even the apparently less efficient design enabled a successful pilot screen when applied to a panel of 21 test genes, since the screen revealed new gene functions in oocysts, overcoming a current roadblock to screening approaches.

In the pilot screen, male fertility phenotypes from our earlier fertility screens^10^ were reproduced perfectly and independently of homing which was expected since mutagenesis with a *Plasmo*GEM-gRNA vector was performed on the male parent. Where the zygote inherits a gene product (mRNA or protein) from the female gamete, our screen did not reveal post-fertilisation functions. The was also expected in the case of *nek4*, because this protein kinase regulates specifically the zygotic cell cycle and is inherited from the female gamete^33^. In contrast, the absence of a more marked homing effect the beta subunit of the mitochondrial ATP synthase, another known female fertility gene^10^, warrants further investigation. It raises the possibility that oxidative phosphorylation is more important in the early mosquito stages than in the growing oocyst, which is relevant in light of efforts to target this pathway for transmission-blocking interventions^34,35^. However, as with other CRISPR screens, absence of a phenotype needs to be corroborated with additional gRNAs or by combining results from different members of a protein complex or enzymes of the same pathway.

The pilot screen revealed a role in oocyst development for a putative transporter, CRTL, which structural modelling showed to be a close homolog of *Pf*CRT. CRTL was investigated previously as a putative apicoplast protein, but was not pursued further when found to be absent from asexual blood stages^36^. We now localise an endogenously tagged protein to the oocyst’s haemozoin crystals, where the most likely location of a protein with multiple transmembrane domains is in the DV, also consistent with the localisation of CRT in blood stages. CRTL is clearly not part of the apicoplast or mitochondria, which have a different distribution in the sporulating oocyst^37,38^ and get incorporated into sporozoites. No oocyst DV marker has to our knowledge been identified previously, and the function of this organelle in nutrient acquisition is unknown. Blood stage parasites ingest haemoglobin from the host cell cytoplasm and degrade it proteolytically it in the acidic environment of the DV. The prosthetic group, heme (ferroprotoporphyrin IX), is oxidized to ferriprotoporphyrin IX (FP) and detoxified by sequestration as microcrystals of hemozoin, a process which is targeted by CQ and other quinoline antimalarials^39,40^. Oocysts inherit the hemozoin crystals from the macrogametocyte, and it is tempting to speculate that a main function of their DVs may be to prevent the release of toxic heme from the hemozoin crystals contained within them. Lysotracker staining shows oocyst DVs are acidified, which would favour the self-association of heme^39^. Deacidified DVs in the CRTL mutant could thus result in the release of toxic heme, which may prevent further oocyst growth.

How CRTL contributes to the acification of the DV remains to be investigated and may well be related to its unknown substrate specificity, which is likely different from that of CRT. *Pf*CRT has been shown to export the doubly-charged chloroquine ion (CQ^2+^) using the outwardly-directed H^+^-gradient in the asexual blood stage DV^30^. The available *Pf*CRT structure was captured with a cavity open to the digestive vacuole, revealing a highly negatively-charged pocket for binding CQ^2+^ (ref. ^30^). Although the natural substrate for *Pf*CRT has been proposed to include host-derived peptides of 4-11 residues^41^, the uptake kinetics in *Xenopus* oocytes is slow and yet to be validated using purified components. Similar to bacterial homologues like YddG, it has been also shown that the purified *Pf*CRT reconstituted into liposomes can export positively-charged amino acids like arginine, with transport rates that seem more consistent with a physiological substrate^30^. Unlike *Pf*CRT, however, the cavity for *Pb*CRTL has both non-polar and positively-charged regions, indicating its substrate is rather a negatively-charged metabolite^42^, possibly with a hydrophobic extension, e.g. ATP (Fig. 6C). While more work is needed to understand the function of CRTL, our data demonstrate how the scalable new screening strategy can lead to the systematic discovery of essential malaria transmission genes whose first essential functions are after fertilisation, thus enabling their potential as targets for transmission-blocking interventions to be assessed.

## Resource availability

### Lead contact

Further information and resource requests should be directed to the lead contact, Oliver Billker.

## Materials availability

Parasite lines and vectors produced by this study are accessible under a material transfer agreement for not-for-profit research purposes. Requests should be made through the lead contact, Oliver Billker.

## Data and code availability

The raw array data from this study have been deposited in github.

## EXPERIMENTAL MODELS DETAILS

### Murine model

Female BALB/c mice were 6 weeks old, and female Wistar Han IGS rats were 150 g upon purchase from Charles River Europe. Mice were group-housed as four cage companions, and rats as two cage companions, in individually ventilated cages with autoclaved chow, sterile drinking water, and sterile wood chip bedding and paper towels for nesting. Environmental conditions were maintained at 21°C, and 55% relative humidity under a 12/12 h light-dark cycle. The animals were acclimatised for 1 week in our animal facility before the experiments. All procedures were performed according to the guidelines and approved protocols from the Umeå Centre for Comparative Biology (UCCB) at Umeå University under Ethics Permit A13-2019 and approved by the Swedish Board of Agriculture.

### Mosquito model

*Anopheles stephensi* mosquitoes were reared and maintained at 28°C with 80% humidity under a 12/12 h light-dark cycle. After being infected with P. berghei parasites, the mosquitoes were kept at 19°C and 80% humidity. Throughout the study, all mosquitoes were provided with an 8% fructose solution (supplemented with PABA and Methyl-4-hydroxy-benzoate) and were anaesthetised using CO2 gas or on ice before dissection.

### Parasite model

*P.berghei* lines used in this study are the wild type reference clone cl15cy1 of *P. berghei* ANKA, the Pb Bergreen line, which expresses GFP from a silent intergenic locus on chromosome 6 under the control of the *hsp70* promoter^43^ and the Pb Bern mCherry line, which uses the same promoter to express mCherry from the p230p locus^44^.

## Method details

### Generation of marker-free single-sex lines expressing BFP-tagged Cas9

Fluorescence and drug resistance marker-free single-sex lines were generated by disrupting PBANKA_145480 (*fd1*) or PBANKA_010240 (*md4*), respectively^6^ using a CRISPR-RGR strategy (ribozyme guide ribozyme) with two *sgRNAs* to target the gene of interest^15^(Fig. S1a) in the *P. berghei* clone cl15cy1. To construct a plasmid to express a CRISPR-Cas9 transgene, we modified the existing CRISPR-RGR strategy with two single guide RNAs (sgRNAs) to target the gene of interest^15^.The sgRNAs were designed using Benchling, and the sgRNA cassette (containing the gRNA along with RGR) was synthesised (Azenta Life Sciences). Subsequently, the sgRNA cassette was cloned into the cas9 plasmid (MH_046) which carries a *hdhfr* selection cassette to generate PL_HC_014 (gRNA plasmid targeting md4) and PL_HC015 (gRNA plasmid targeting *fd1*; Table S4, S5). DNA fragments encoding an *hsp70* promoter, N-terminal flag tagged *cas9*, *bfp* and *dhfr* 3’UTR were cloned into a plasmid and flanked with 500 bp sequences targeting either *fd1* or *md4* locus (Fig. S1a).

To generate the repair template, *hsp70* promoter, *cas9, bfp* and *Pbdhfr* 3’UTR gene sequences were amplified from plasmids Pb_MH21 and R6K-GW-BFP (PL_HC_001) respectively using Advantage 2 Polymerase Mix (TaKaRa). Individual fragments were assembled by stitch PCR, digested with *Kpn*1 and *Sac*11, and ligated into an intermediated vector (pMisc 017) to generate PL_HC013. The *cas9-bfp* fragment was sub-cloned into two pre-synthesized plasmids (Azenta Life Science) containing 500 bp sequences upstream and downstream of either *fd1* or *md4* gene locus to generate plasmid carrying repair template PL_HC_018 and PL_HC_017 respectively.

To generate transgenic single-sex lines expressing Cas9, *P. berghei* schizonts were co-transfected with 1 μg of the Cas9-sgRNA vector and linearized cas9bfp repair template targeting either the *md4* or *fd1* gene locus. For each transfection, 1 μl of isolated schizonts was mixed with 7 μl of DNA and 18 μl of P3 Primary Cell 4D-Nucleofector solution from Lonza. This mixture was then added to a well within a 4D-Nucleofector 16-well strip from Lonza and electroporated using the FI-115 program on the Amaxa Nucleofector 4D. Transfected parasites were promptly injected intravenously into BALB/c mice and were subjected to selection with 0.07 mg/mL of pyrimethamine in drinking water starting from day one post-infection. Pyrimethamine selection was terminated on Day 5, and mice were administered 5-fluorocytosine (1 mg/mL, Sigma) via drinking water to eliminate the CRISPR plasmid carrying the *dhfr/yfcu* selection cassette (negative selection). Insertion of *cas9 bfp* in the *md4* and *fd1* gene loci was confirmed by PCR. Cas9-BFP expression in the nucleus was visualised by live imaging of infected cells in the Zeiss Axio imager 2 fluorescent microscope. The images were analysed using Fiji.

### FACS sorting and dilution cloning of single-sex lines

To generate clonal lines of Cas9-expressing *fd1^-^* and *md4^-^* parasites, infected mice were euthanized when parasitemia reached 0.1%. Blood (10 μl each) was collected in 200 μl of CLB buffer (PBS, 20 mM HEPES, 20 mM Glucose, 4 mM sodium bicarbonate, 1 mM EDTA, 0.1% w/v bovine serum albumin, pH 7.25). Erythrocytes were then selected by gating on forward/side-light scatter, and BFP-expressing parasites were gated in the BV421A channel using the BD FACS Aria III. These BFP-positive parasites were sorted into cold CLB and intravenously injected into BALB/c mice. Each clonal line was re-genotyped to verify modifications in the *md4* and *fd1* gene loci and to ensure the absence of the *dhfr*-resistance cassette (*Cas9* plasmid).

### Generation and genetic crossing of myoA tagged lines

MyoA tagging vectors were engineered in the *Plasmo*GEM *myo*A gDNA library clone pbG01_2365g05 using lambda Red-ET recombinase-mediated engineering in *E.coli*^14^. In short, two sets of recombineering oligos carrying *attR1*, *attR2*, and barcode were designed that have 50 bp homology to the 3’ UTR of *myoA* gene. The second set of recombineering oligos had a recognized section within the forward primer to incorporate a shield mutation in the final tagging vector. These primers were employed to amplify the *zeo-PheS* cassette from the *pR6K-attL-zeo-phes-attR2* plasmid. The resulting PCR products were then utilised for recombineering to generate the intermediate vectors PL_HC_020 and PL_HC_021. PL_HC_020 harbors the zeo-phes selection integrated into the C-terminus of the myoA in the library clone, while PL_HC_021 contains a shield mutation at the 3’ end of the myoA gene and a portion of the 3’UTR removed to protect the gene from cleavage by gRNA 1 and 2, respectively. The clones were positively selected using zeocin and were sequenced. These intermediate vectors were further modified into final transfection vectors using a Gateway recombinase reaction, integrating the *dhfr* resistance cassette into the vector. The mCherry tagging vector was produced by performing a gateway recombinase reaction between PL_HC_020 and the *Plasmo*GEM *pR6K_mcherry* gateway plasmid, generating PL-HC_022 (Supplementary Fig 2b). Two variants of myoA GFP tagging vectors were generated. The first one (PL-HC_028) was produced by a recombinase reaction between the *Plasmo*GEM pR6K gfp gateway plasmid and PL_HC_021. The second GFP tagging vector (PL_HC_026) was generated by a recombinase reaction between a modified *Plasmo*GEM vector carrying a gRNA cassette targeting the myoA gene and 3’UTR with PL_HC_026 (Supplementary Fig 2a&b). These vectors were used to transfect *fd1^-^cas9* and *md4^-^cas9* parasites, following the procedures outlined earlier. Transfected parasites were selected on pyrimethamine and were dilution cloned. Each clonal line was genotyped to confirm modifications in the *myoA* gene loci. Genetic crosses between these lines were conducted by infecting mice with various combinations of *myoA-*tagged male and female-only parasites at a 1:1 ratio. Infected mice were anaesthetised and utilised for the direct feeding of *A. stephensi* mosquitoes.

### DNA extraction for genotyping

To extract the parasite from the host blood, heparinized whole blood was collected by cardiac puncture when the parasitemia was approximately 3%. To remove blood plasma, 100 μl of whole blood was pelleted by centrifugation for 8 minutes at 450g. The cells were then resuspended in 1 ml of RBC lysis buffer (150 mM NH4Cl, 10 mM KHCO3, 1 mM EDTA) and incubated on ice for 15 minutes. After lysis, parasites were pelleted at 450 x g for 8 minutes and washed 2 times with 1X PBS, or until complete removal of red pigment of blood. The parasite pellets were either stored at - 80°C or DNA from these parasite pellets was isolated using Thermo Scientific GeneJET Genomic DNA Purification Kit according to manufacturer instructions, and genotyping was performed using advantage 2 polymerase mix (TaKaRa). The genotyping strategies are described in supplementary figures 1, 2 and 4. The primers used for genotyping are mentioned in Supplementary Table S4.

### Western blot assay

Parasite pellets (generated as described for DNA extraction) were resuspended in 10 μl of parasite lysis buffer (SDS 4%, 0.5 % Triton X-100, 0.5x PBS). The lysate was combined with 4X Laemmli loading buffer supplemented with 20% β-mercaptoethanol. Samples were boiled at 95 °C for 5 minutes and loaded onto a 4-20% Bio-Rad TGX precast gel. Electrophoresis was performed at 90 V for 2 h. Proteins were transferred onto a PVDF membrane in the Turbo Transblot (Bio-Rad) using the High Molecular weight program (10 minutes, 2.5 A constant, up to 25 V). The membrane was blocked with 2%BSA overnight and was probed with antibodies against goat anti-FLAG (1:1000, Cell Signaling Technologies, D6W5B, #14793), alpha Tubulin (1:10000, mAb, mouse) or rabbit anti-HA (1:1000, Cell Signaling Technologies) followed by a compatible HRP-linked secondary antibody. The blots were developed using Immobilon Western Chemiluminescent HRP Substrate (Merck-Millipore) and were visualised on an Amersham Imager 680.

### Exflagellation and ookinete conversion assay

BALB/c mice were intraperitoneally injected with 200 μl of phenylhydrazine (6 mg/ml; Sigma). Three days after treatment, mice were infected with *P. berghei* parasites. Blood from infected mice exhibiting 8-10% gametocytaemia was added to ookinete medium (20% FBS, 100 μM xanthurenic acid (Sigma), 24mM sodium bicarbonate, and RPMI (Gibco 52400-025), pH 8.2). For the exflagellation assay, infected blood was added at a 1:5 ratio of blood to ookinete medium, and parasites were incubated for 10 minutes. Male exflagellation events were counted in a standard hemocytometer under a light microscope. To determine ookinete conversion rates, infected blood was added to ookinete media at a 1:5 ratio and incubated for 23 hours at 19°C to allow for ookinete formation. After incubation, 1 ml of ookinete culture was taken and mixed with 1 ml of 4% paraformaldehyde (PFA) and incubated at room temperature for 15 minutes. Fixed ookinetes were pelleted at 500xg for 3.5 minutes and washed twice with 1X PBS. The final pellet was resuspended in 100 μl of PBS. From the resuspension of the fixed ookinete, 15 μl was mixed with 50 μl of staining solution (Cy3-labelled 13.1 mouse monoclonal anti-P28 (1:500 dilution) and Hoechst (1:2000)). The samples were incubated at room temperature (RT) for approximately 12 minutes. 6 μl of the stained sample was placed on a glass slide and covered with a vaseline-edged coverslip. The samples were visualised in a Zeiss Axio Imager 2 fluorescent microscope. The images were analysed using Fiji. The number of ookinetes (banana-shaped cells that are Cy3-positive) and female gametocytes (round cells that are Cy3-positive) were quantified. The conversion rate was calculated as the percentage of Cy3-positive ookinetes to Cy3-positive macrogametes and ookinetes.

### Indirect Immunofluorescence

Blood-stage parasites and ookinetes were fixed in 4% PFA in PBS for 40 and 15 minutes respectively at RT. Fixed cells were washed with 1XPBS and were resuspended in PBS. Cells were added to poly-D-lysine coated glass slides and were allowed to settle for 15 mins. Permeabilization of the parasites was done using 0.2% v/v Triton X-100 in PBS for 5 min at RT. Permeabilized parasites were washed with 1X PBS and then blocked with 3% w/v bovine serum albumin (BSA) in PBS for 40 minutes at RT. Parasites were stained overnight with 13.1 mice monoclonal anti-P28 (1:1000) and rabbit anti-HA (1:250, Cell Signaling Technologies) antibodies. Parasites were washed 3 times in PBS for 10 min each and then were stained with secondary antibodies (Thermo Fischer) anti-mouse Alexa Fluor 647 (1:500) and anti-rabbit Alexa Fluor 488 (1:500) for 2h. The cells were washed and stained with Syto 19 or Hoechst to visualise nuclear DNA. Images were acquired using a Leica SP8 confocal microscope and were visualised using Fiji. For live cell imaging of blood and ookinete stage parasites, cells were stained with LysoTracker Deep Red (1 μM, Invitrogen) or LysoTracker Green DND-26 (1 μM, Invitrogen) for 10 minutes at RT and were counterstained with Hoechst to visualise DNA. 6 μl of the stained sample was placed on a glass slide and covered with a vaseline-edged coverslip. The samples were visualised in a Zeiss Axio Imager 2 fluorescent microscope. The images were analysed using Fiji.

### *P.berghei* mosquito infection and transmission

Mosquitoes were fed on mice infected with *P.berghei* at a parasitemia of 3-5%. To determine oocyst number and size, midguts were dissected on the indicated day post-infection and visualised using a Zeiss Axio Imager 2 fluorescent microscope. Oocyst numbers and size were quantified using built-in plugins in Fiji. Localization of the HA-tagged protein in the oocyst was performed as previously described^45^. Briefly, infected mosquito midguts were fixed in 4% PFA containing 0.1% saponin in PBS and incubated for 45 minutes on ice. After three washes with PBS/0.1% saponin of 15 minutes each, parasites were blocked with 3% BSA, 0.1% saponin in PBS for 30 minutes on a thermal shaker at room temperature (RT). Parasites were stained overnight with rabbit anti-HA (1:200, Cell Signaling Technologies) and then stained with anti-rabbit Alexa Fluor 488 (1:500, Thermo Fisher Scientific) for 2 hours. Cell nuclei were labelled with Hoechst. For live imaging of oocysts and visualisation of acidic compartments in the oocyst, the midgut was dissected into PBS containing LysoTracker™ Deep Red (1 μM, Invitrogen) or LysoTracker Green DND-26 (1 μM, Invitrogen) along with Hoechst and incubated for 10 minutes at RT before imaging with a Zeiss Axio Imager 2 fluorescent microscope. Time-lapse images of the oocysts were captured by imaging live oocysts over a period of 30 seconds in Zeiss Axio Imager 2. Salivary gland sporozoites in the infected mosquitoes were imaged on day 18 and day 20. To assess mosquito-to-mouse transmission, approximately 15-20 infected *A.stephensi* mosquitoes were fed on 3 anaesthetised BALB/c mice. Mouse parasitemia was monitored until day 8 post-mosquito bite by Giemsa staining.

### Generation of homing vectors

Homing vectors were generated by recombineering the gRNA cassette to existing *Plasmo*GEM vectors. To do so, we generated a plasmid by Gibson assembly to generate an MCS (multiple cloning site) sandwiched between the *Pb hsp70* promoter and the *P. yoelii dhfr* terminator (PL_HC_030). The plasmid also had a zeocin resistance gene cloned adjacent to the *Pb hsp70* promoter. The ribozyme guide ribozyme (RGR) sequences containing two sgRNA targeting the gene of interest were synthesised (Azenta life science) and were cloned into the MCS of PL_HC_030. The zeocin resistance gene along with the *hsp70* promoter, gRNA cassette, and *dhfr* terminator was amplified with primers containing homology regions (50 bp each) to either side of the 3xHA part of the *Plasmo*GEM KO vector. The resulting PCR products were then utilised for recombineering into *Plasmo*GEM vectors to generate homing vectors.

### Generation of mutant pools using *Plasmo*GEM and homing vectors

To generate pools of *Plasmo*GEM and homing vectors, 21 transformed *E.coli* TSA cells carrying knock-out vectors were inoculated into 96 well plates containing 1 ml of Terrific Broth with kanamycin (30 μg/ml) and were grown overnight at 37°C. The cultures were then pooled and plasmid was extracted using the QIAGEN Plus Midi Prep Kit. A total of 30 μg (approximately 1 μg of each vector) was digested overnight with NotI to release the targeting vector from the linear plasmid backbone and was used for three independent transfections as described previously^9^. Briefly, *fd1^-^cas9* schizonts derived from infected rats were harvested after 22 hr culture and purified on a Nycodenz (Sigma) gradient. Purified schizonts were electroporated with either *Plasmo*GEM or a homing vector pool using the FI115 program on the Amaxa Nucleofector 4D (Lonza). Transfected parasites were then injected intravenously into BALB/c mice, and were selected with pyrimethamine (0.07mg/mL) in drinking water from day one post-infection.

### Transmission of barcoded KO parasites through mosquitoes

After seven days post-transfection with the *Plasmo*GEM or homing vector pool, mice with a parasitemia of 3-5% were euthanized and blood was collected via cardiac puncture. The blood from *fd1^-^cas9* mutant parasites was mixed with a female donor line to establish a final ratio of infected red blood cells between these parasites at 1:1. This mixture was used to infect BALB/c mice. Mice with a parasitemia of 3-5% on day 3-4 post infection were selected to feed *A. stephensi* mosquitoes. For each biological replicate, 80-100 infected mosquito midguts were dissected for genomic DNA extraction. The female donor line used was GIMO *cas9*, which is capable of selfing but carries no barcode module.

### Library preparation, barcoded sequencing, and analysis

DNA was extracted from three different samples. The vector pool DNA was extracted from the left-over media in the transfection cuvette (Barcode input 1). This served as input control and was used to quantify the initial diversity of the barcodes/vectors. Parasite genomic DNA was extracted from the mouse blood sampled after feeding the mosquitoes (Barcode input 2) using DNeasy Blood and Tissue Kit (Qiagen). The parasite gDNA was extracted from the infected mosquito midguts 14 days post-infection (Barcode output) by boiling the midguts in Quick Extract DNA Extraction Solution (Epicentre). *Plasmo*GEM barcodes were amplified by PCR in using the extracted DNA samples and the primers 91_Illumina and 97_Illumina (Table S6). The PCR amplicons were used as an input for a second PCR to add sample-specific index tags using primers shown in the Table S6. The resulting libraries were then pooled and sequenced using the Illumina MiSeq Reagent Kit v2 (300 cycles), which were loaded at a low cluster density (4×105 clusters/mm2) with 50 % PhiX.

The index tags were utilised to segregate sequencing reads for each sample. The total number of barcodes for each gene within the sample was counted using a Python script. Relative abundance was calculated for each mutant by dividing the mutant barcode count by the total number of barcodes in the sample. The fold change in relative abundance between the mutant barcode in the mosquito midgut (barcode output) and that in the mouse blood (barcode input 2) was calculated and normalised with spike-in control barcodes for genes whose knockout did not exhibit a phenotype in the mosquito stages (KIN, PBLP, and PBANKA_0308200) to generate the oocyst conversion rate. The fold change in relative abundance between the mutant barcode in the mouse blood (barcode input 2) and that in the transfection cuvette (barcode input 1) was used to calculate the off-target effect of gRNA for each gene (Fig 3A). Barcode counts for each gene can be found in Table S3. The details of the analysis are deposited in GitHub.

### Generation of CRTL KO and tagged transgenic lines

The *Plasmo*GEM knock out and tagging vectors PbGEM-93381and PbGEM-93389 were used to generate *crtl^-^* and *crtl*-3xHA tagged parasites in Pb Bern mCherry and *P. berghei* ANKA cl15cy1 background respectively. Briefly, *P.berghei* schizonts were electroporated with 3 mg of each plasmid using the FI115 program on the Amaxa Nucleofector 4D (Lonza). Transfected parasites were promptly injected intravenously into BALB/c mice and were subjected to selection with 0.07 mg/mL of pyrimethamine in drinking water starting from day one post-infection. Clonal lines were derived by limited dilution cloning and were genotyped to verify modifications in the PBANKA_0916000 gene locus.

### TEM

Mosquito midgut was fixed with 2.5% Glutaraldehyde (TAAB Laboratories, Aldermaston, England) in 0.1 M phosphate buffer. Samples were further post-fixed in 1% aqueous osmium tetroxide dehydrated with ethanol and finally embedded in Spurr’s resin (TAAB Laboratories, Aldermaston, England). All steps were performed using the Pelco Biowave pro+ (Ted Pella, Redding, CA). 70 nm ultrathin sections were picked up on copper grids and post-stained with 5% aqueous uranyl acetate and Reynolds lead citrate. Grids were examined with Talos L120C (FEI, Eindhoven, The Netherlands) operating at 120 kV. Micrographs were acquired with a Ceta 16M CCD camera (FEI, Eindhoven, The Netherlands) using Velox ver 2.14.2.40.

## AUTHOR CONTRIBUTIONS

Conceptualization, O.B and A.B.; Methodology, O.B, A.B and M.H.; Software, V.P.; Formal Analysis, O.B, A.B., M.H, V.P., D.D., Investigation, A.B., M.H., P.T., D.D.; Writing – Original Draft, A.B.; Writing – Review & Editing, O.B, M.H., D.D.; Visualization, O.B., A.B., M.H, V.P., D.D.; Supervision, O.B., A.B; Project Administration, A.B.

## Acknowledgment

The authors are grateful to Sarah Lundgren, Divya Das, and Rashmi Mishra, who bred and prepared the mosquitoes for this work. SciLifeLab National Genomics Infrastructure in Stockholm is acknowledged for sequencing support. The authors acknowledge Chiara Currà, Inga Siden-Kiamos (Foundation for Research and Technology— Hellas, Institute of Molecular Biology and Biotechnology), and Rita Tewari (University of Nottingham) for the valuable inputs. The authors acknowledge the facilities and technical assistance of the Umeå Core Facility Electron Microscopy (UCEM) at the Chemical Biological Centre (KBC), Umeå University, a part of the National Microscopy Infrastructure NMI (VR-RFI 2019-00217).

