## Supplemental Figures S1-S8 for "A CRISPR homing screen finds a chloroquine resistance transporter-like protein of the *Plasmodium* oocyst essential for mosquito transmission of malaria"

### Figure S1

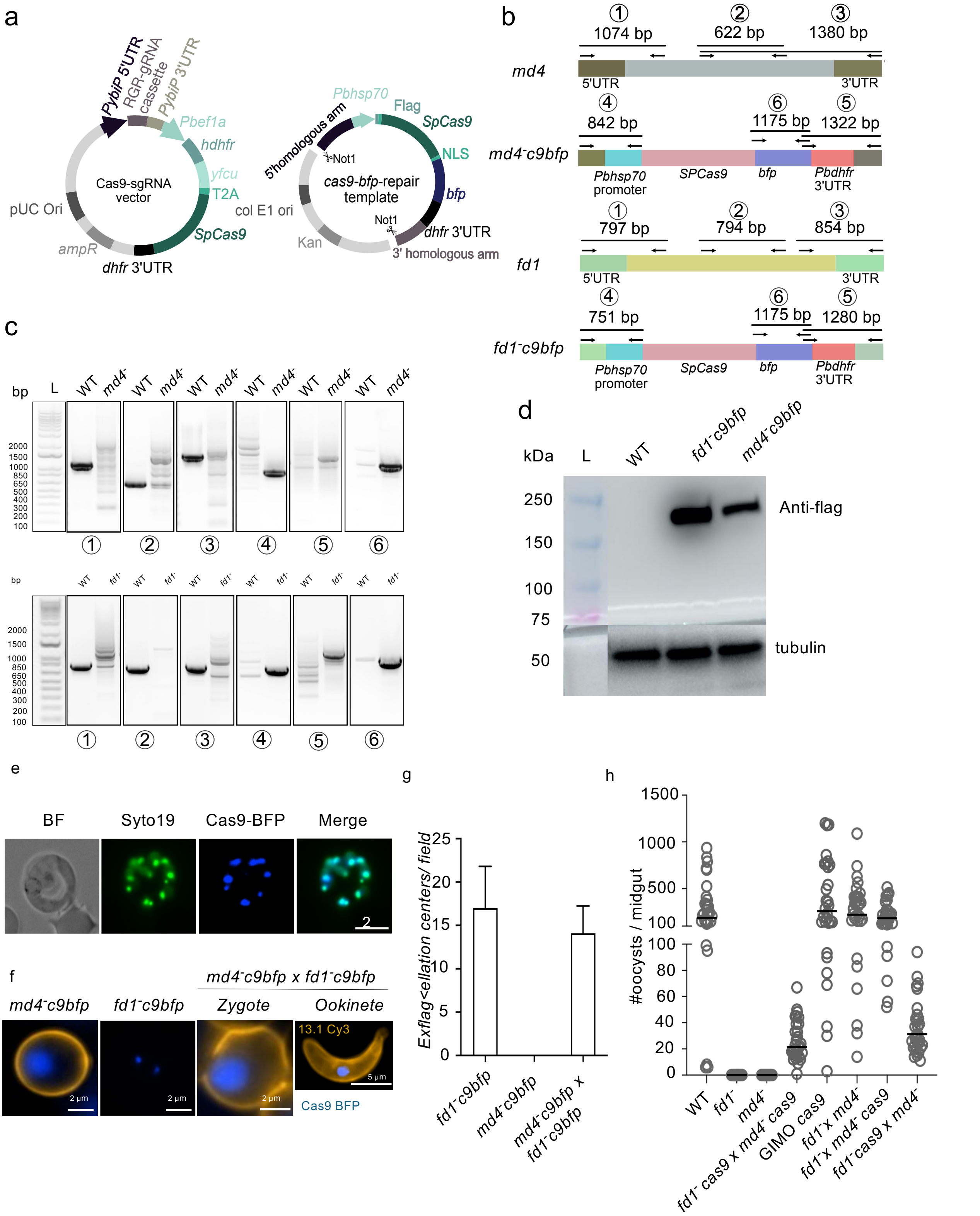

**Supplementary Figure 1: Generation of *Cas9-bfp expressing* single sex lines. (a)** Schematic of vectors used to generate single sex lines. **(b)** Schematic showing primers used to genotype *cas9-bfp* lines. **(c)** Genotyping of *cas9-bfp* single sex lines. **(d, e)** Nuclear localization and expression of Cas9-BFP assessed by microscopy and western blot in mixed blood stages **(f)** Expression of Cas9 BFP in activated female gametes (*md4⁻c9bfp)*, male gametes (*fd1⁻c9bfp)*, zygote and ookinete (*md4⁻c9bfp x fd1⁻c9bfp)*. **(g)** Assesment of single sex lines to form active male gametes by exflagellation assay. **(h)** Fitness of Cas9-expressing single sex lines determined by counting the number of oocyst in the mosquito midgut 12 days post infectious blood meal. The data are from two independent experiments with 25 infected mosquitoes in each set.

### Figure S2

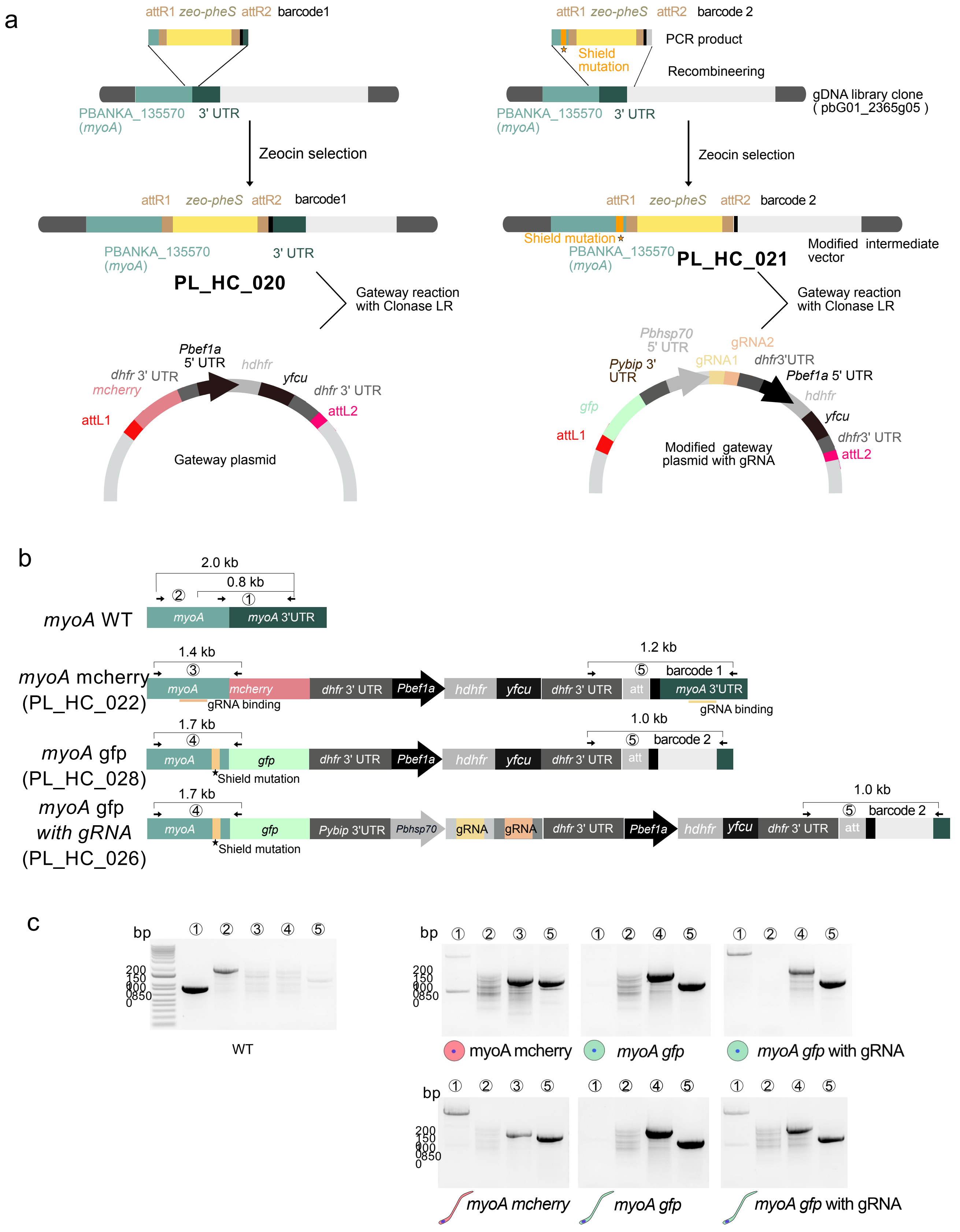


**Supplemental Figure 2.** Generation of *myoA mcherry* and *gfp* tagged parasites expressing gRNA in male (*fd1*⁻) and female (*md4*⁻) *cas9-bfp* lines. **(a)** Shematic showing the generation of modified tagging vectors that was used to make myoA mcherry and gfp tagged paraites expressing gRNA. **(b)** Schematic of final tagging vector **(c)** Genotyping of male (*fd1*⁻) and female (*md4*⁻) -only *cas9* expressing lines transfected with the tagging vectors.

### Figure S3

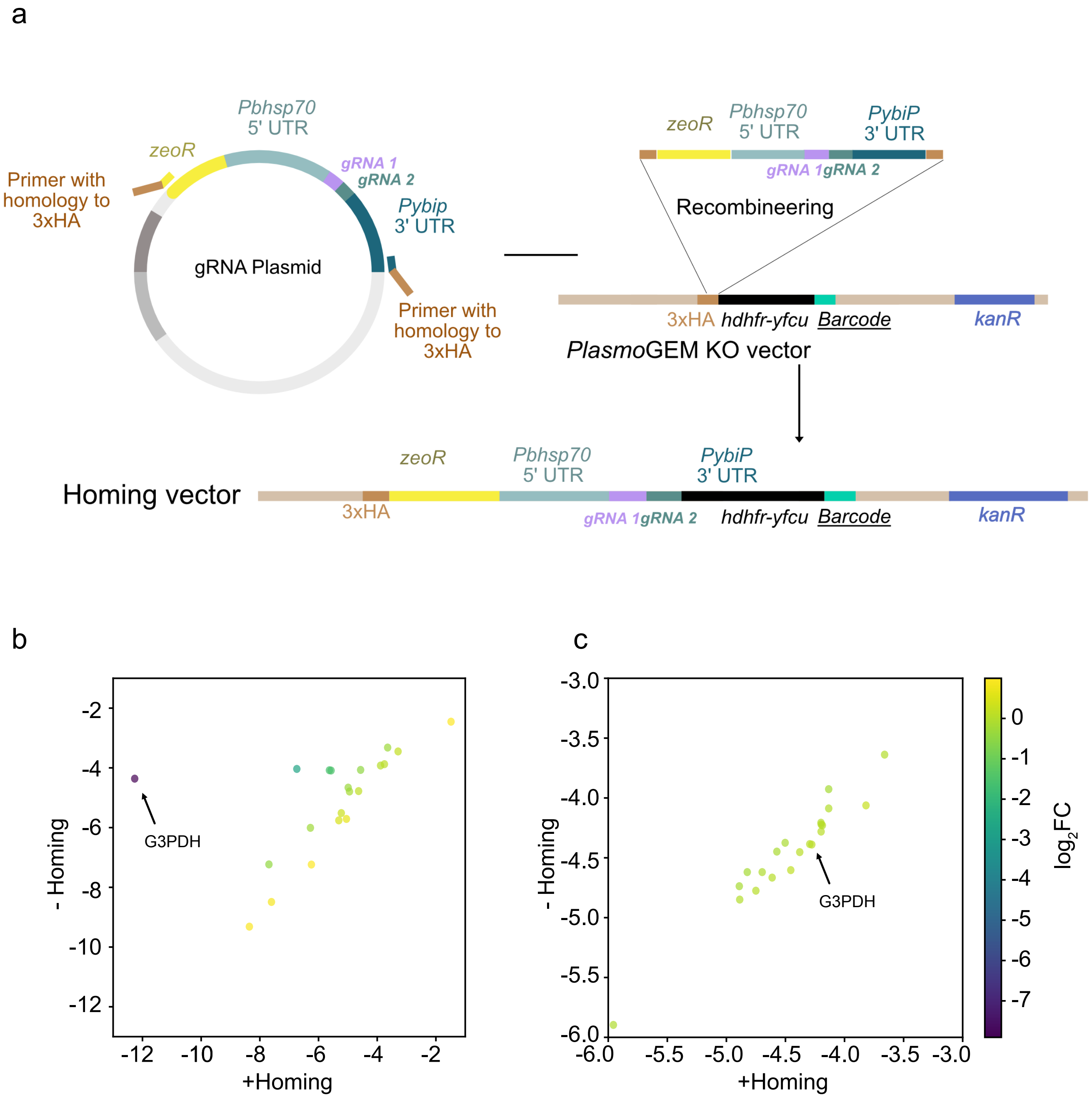


**Supplemental Figure 3. Preparation of homing-Cas9 vectors for the pilot screen and identification of possible G3PDH off-target effect. (a)** Schematic showing the generation of homing vectors for the pilot screen. (b) Off target score of gRNA in homing vectors visualized by a change in relative abundance of the gene in the blood stages when transfected with homing vector (+ homing) in comparison with *Plasmo*GEM vector (-homing). **(c)** Change in relative abundance of the homing vector in the cuvette input (+homing) in comparison with *Plasmo*GEM vector (-homing) is plotted. Different colors indicate difference in $\log_2$-fold change between + homing in comparison to -homing.

### Figure S4

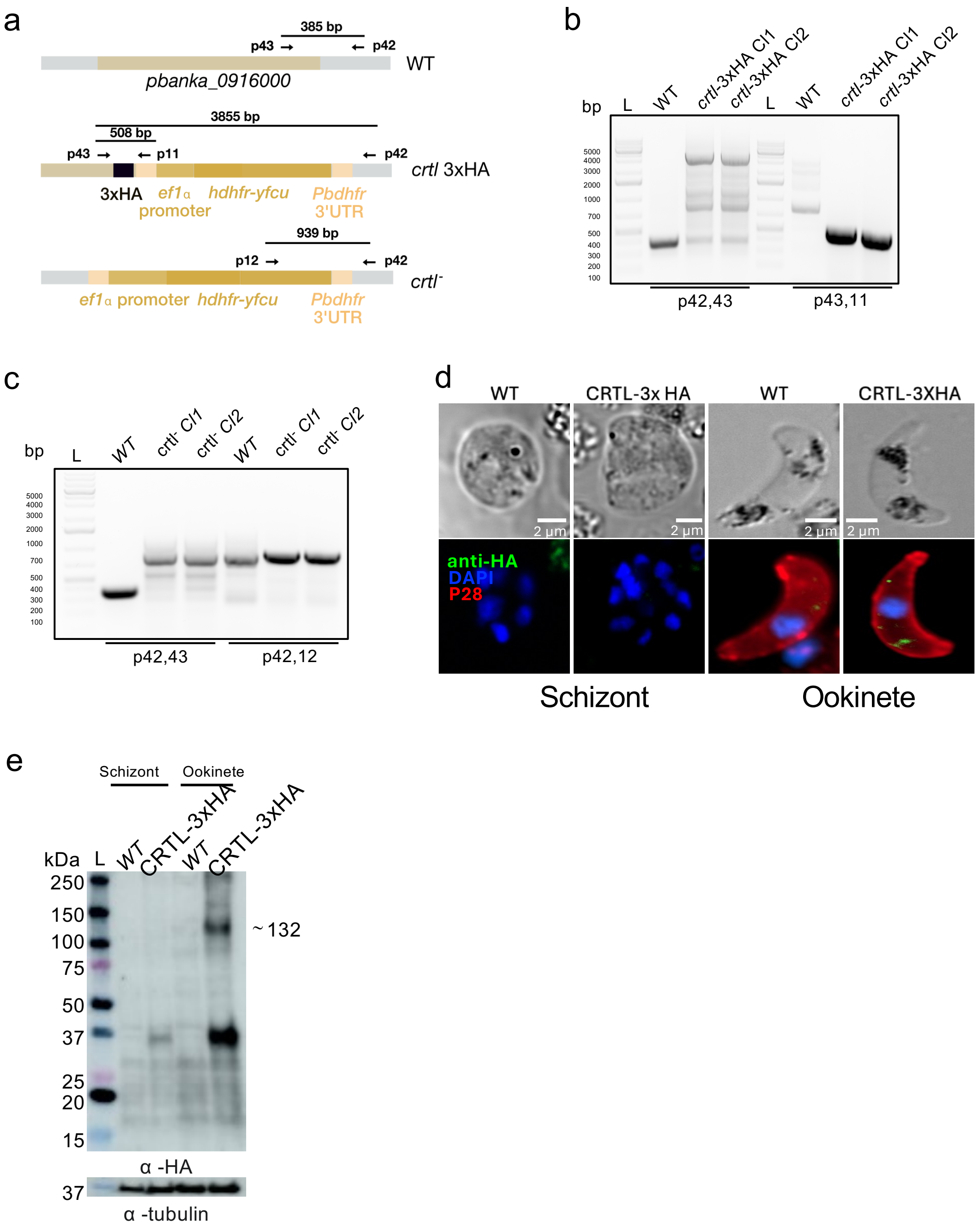


**Supplemental Figure 4.** Generation of *crtl*$^{-}$ and 3X HA tagged lines. **(a)** The Schematic of WT, *crtl*$^{-}$ and crtl 3xHA gene locus. Primers that were used for genotyping are highlighted. **(b, c)** Genotyping of *crtl* 3x-HA tagged and *crtl*$^{-}$ lines. **(d, e)** Localisation and expression of CRTL-3xHA in fixed schizont and ookinetes analysed by fluoroscent microscopy and westren blot respectively. Wild type (WT, untagged) parasites are used as a negative control. Hoechst (blue, DNA); CRTL-HA (green); P28 (red).

### Figure S5

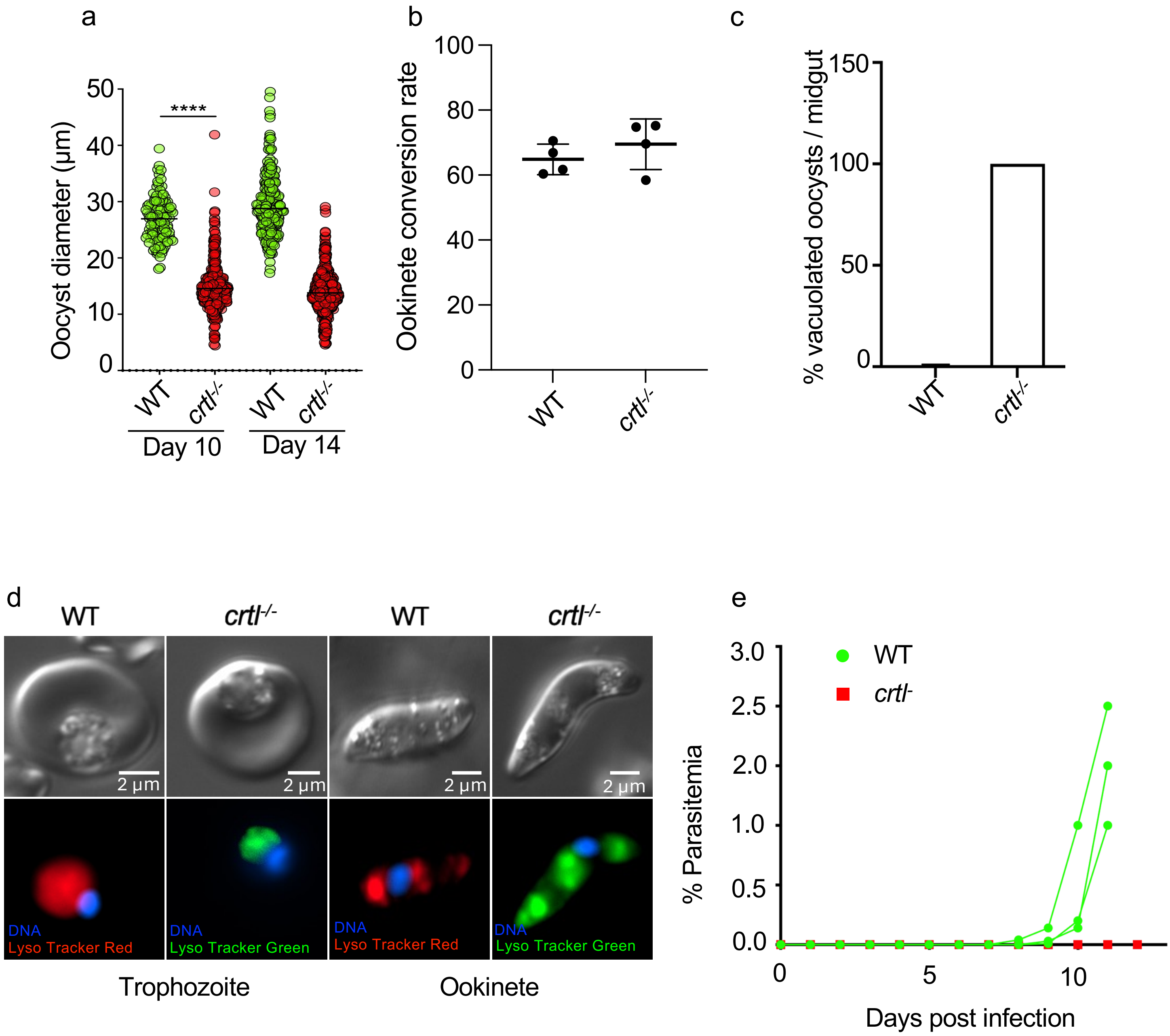


**Supplemental Figure 5.** Characterisation of a *crtl* knockout clone.
(a) The diameter of individual *Pb Bergreen* (WT) or *Pb mcherry crtl* knock out oocysts on day 10 and day 14 post-infection is quantified and plotted. The data are from 2 independent experiments with 25 infected mosquitoes in each set.(unpaired t test ****P < 0.0001, ns not signifnicant). (b) The percentage conversion from female gametocyte to ookinete is quantified and plotted. The data is from 4 independent experiments. (c) Quantification of percentage vacuolated oocyst in the WT and *crtl*⁻ day 10 infected mosquito midgut. The data is from 20 randomly selected infected mosquito midgut. (d) A representative image of *Pb Bergreen* (WT) or *Pb mcherry crtl* knock out blood stage parasites and ookinetes after staining with either LysoTracker Red (WT) or Lyso Tracker Green (*crtl*⁻) to visualize acidic compartments. Hoechst was used to stain DNA. (e) Parasitemia in mice after transmission of WT or *crtl*⁻ parasites by infected mosquito bites (15 infected mosquitoes/ mice). Three mice were used for each infection.

### Figure S6

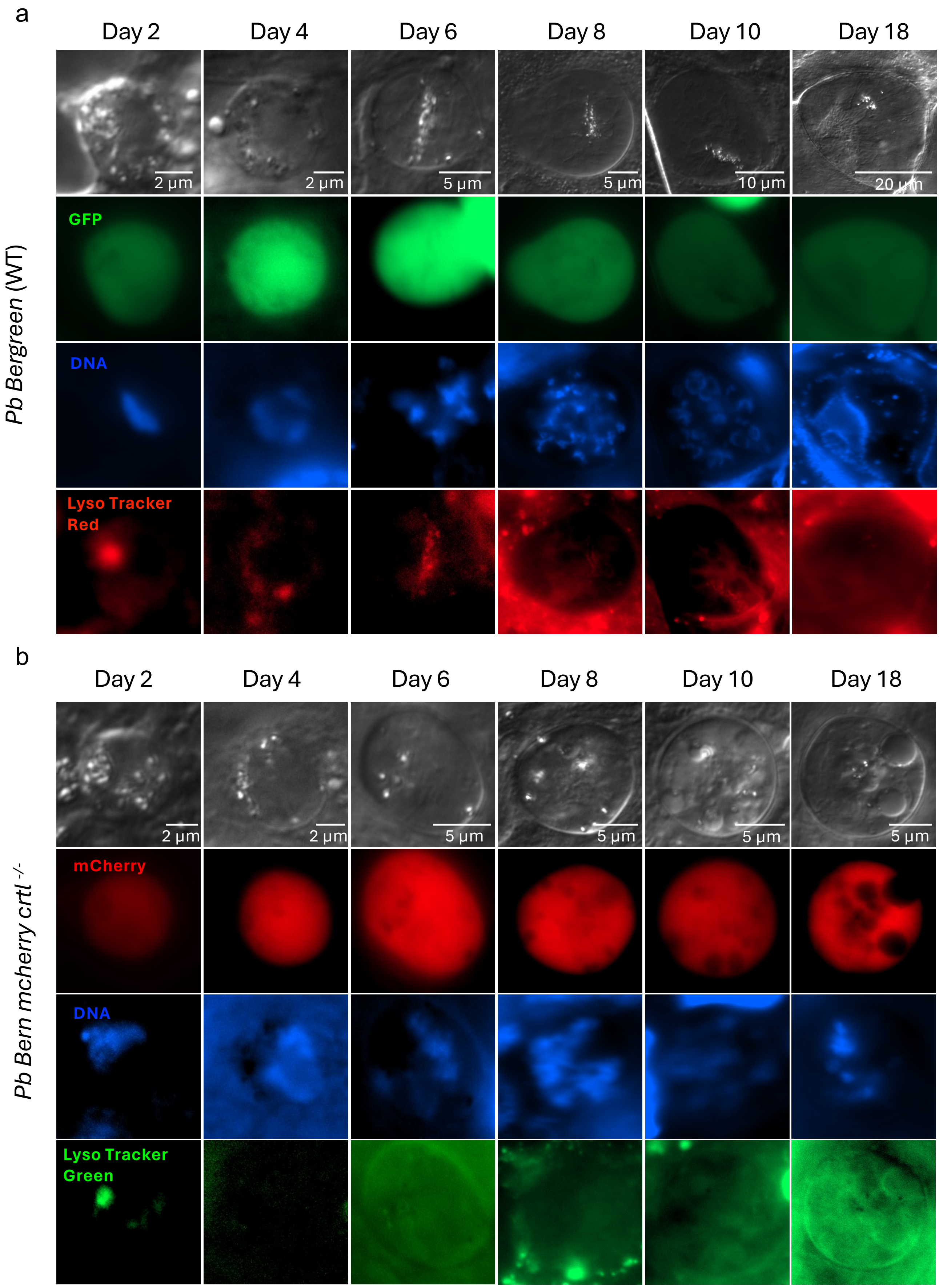


**Supplemental Figure 6.** Fluorescence micrographs showing stages of oocyst development.

(a) Representative images of GFP expressing *P. berghei* oocysts stained with LysoTracker Red and Hoechst for DNA.

(b) Representative images of *mcherry* expressing *crtl*⁻ knock out ocysts stained with Lyso Tracker Green and Hoechst for DNA.

### Figure S7

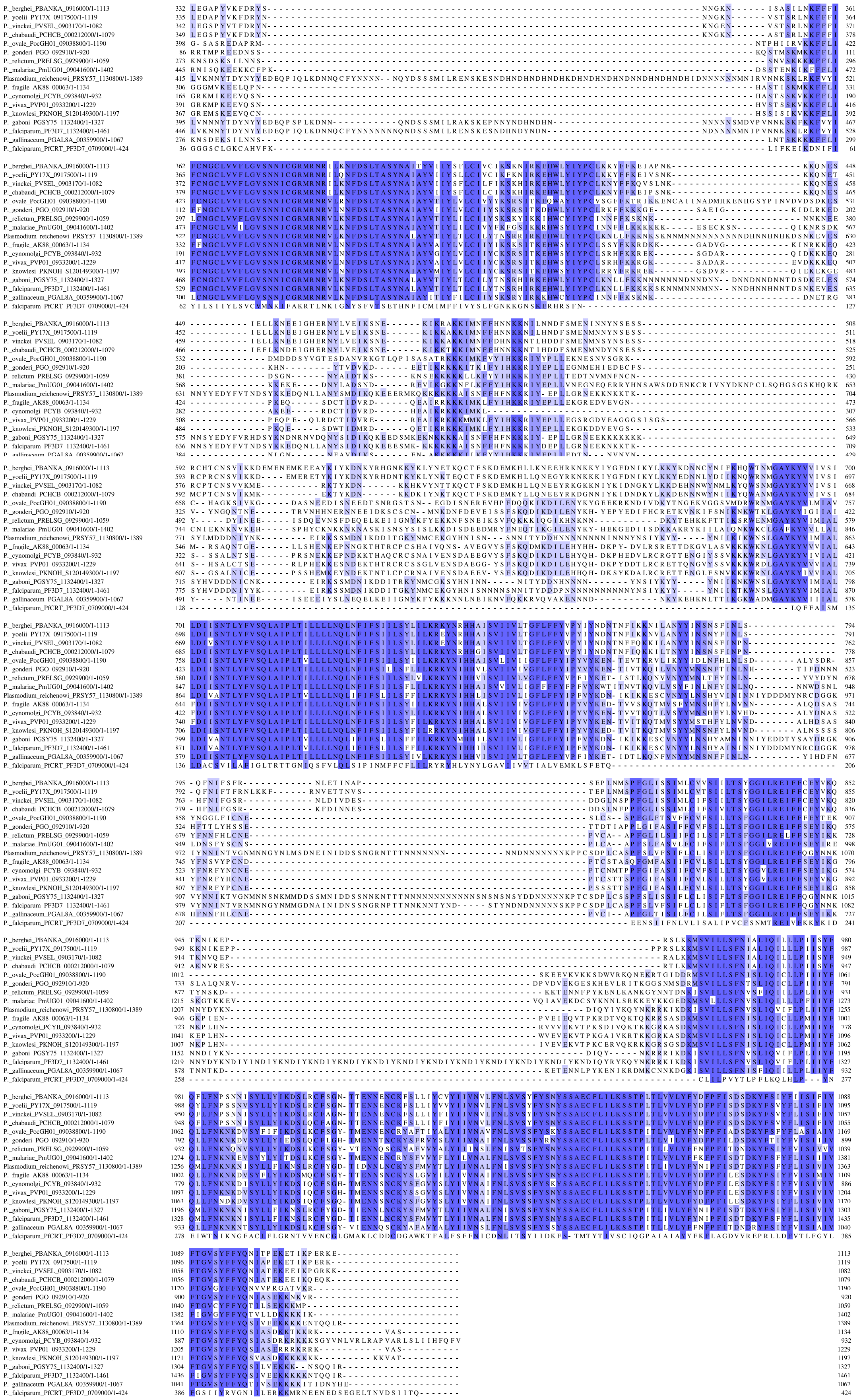

**Supplemental Figure 7.** Multiple sequence alignment of CRTL homologues from different *Plasmodium* species with PfCRT.

### Figure S8

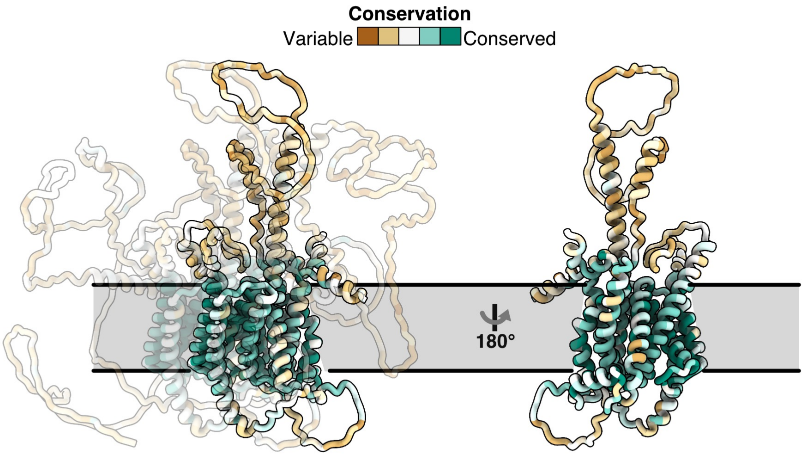


**Supplemental Figure 8.** Two views of the core of *P. berghei* CRTL (as modelled on PfCRT) are shown and coloured according to conservation in the genus *Plasmodium* from brown (not conserved) to teal (high conservation). Conservation of each residue is based on the amino acid alignment in Fig. S7, which uses the following sequences: *P. berghei* PBANKA_0916000, *P. yoelii* PY17X_0917500, *P. vinckei* PVSEL_0903170, *P. chabaudi* PCHCB_000212000, *P. ovale* PocGH01_09038800, *P. gonderi* PGO_092910, *P. relictum* PRELSG_0929900, *P. malariae* PmUG01_09041600, *P. reichenowi* PRSY57 1130800, *P. fragile* AK88_00063, *P. cynomolgi* PCYB_093840, *P. vivax* PVP01_0933200, *P. knowlesi* PKNOH_S120149300, *P. gaboni* PGSY75_1132400, *P. falciparum* PF3D7_1132400, *P. gallinaceum* PGAL8A_00359900 and *P. falciparum* PfCRT_PF3D7_0709000. Only conserved regions are shown.
