## Supplemental Videos for "A CRISPR homing screen finds a chloroquine resistance transporter-like protein of the *Plasmodium* oocyst essential for mosquito transmission of malaria"

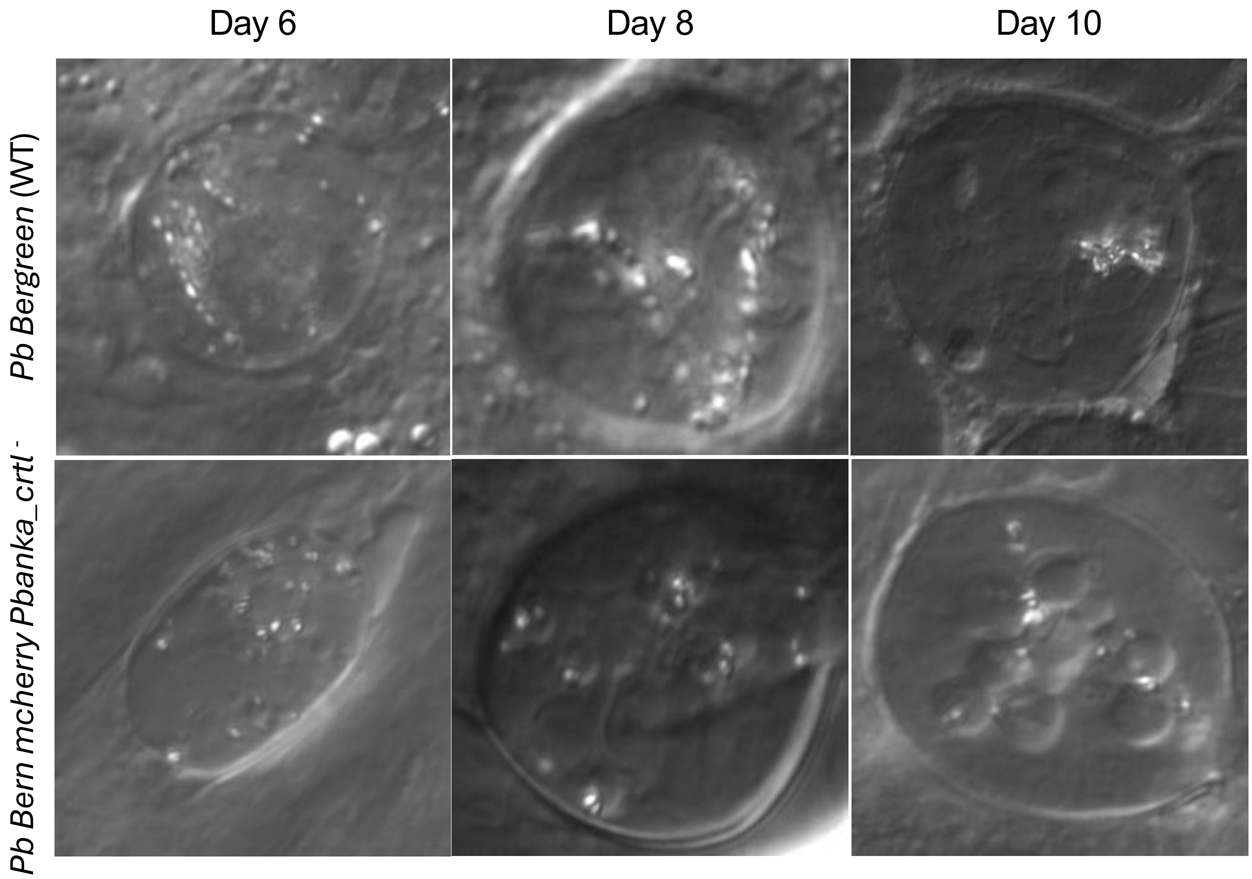


**Supplemental Videos 1. CRTL stabilizes digestive vacuole-like structures and prevents vacuolisation of the oocyst.** Representative timelapse videos of *Plasmodium berghei* oocysts days post infection with either *Pb Bergreen* or *Pb mcherry crtl-* parasites. The digestive vacuole-like dense structures in the oocysts exhibit mobility till day 6. By day 8 in the WT oocyst, these structures become stabilized. *crtl-* oocysts, in contrast, become hyper-vacuolated and vacuoles and hemozoin crystals remain mobile.
